# The growth rate of senile plaques is determined by the competition between the rate of deposition of free Aβ aggregates into plaques and the autocatalytic production of free Aβ aggregates

**DOI:** 10.1101/2024.04.06.588435

**Authors:** Andrey V. Kuznetsov

**Affiliations:** Department of Mechanical and Aerospace Engineering, North Carolina State University, Raleigh, NC 27695-7910, USA

**Keywords:** neuron, Alzheimer’s disease, Finke−Watzky model, mathematical modeling, neurotoxicity, polymerization

## Abstract

The formation of amyloid beta (Aβ) deposits (senile plaques) is one of the hallmarks of Alzheimer’s disease (AD). This study investigates what processes are primarily responsible for their formation. A model is developed to simulate the diffusion of amyloid beta (Aβ) monomers, the production of free Aβ aggregates through nucleation and autocatalytic processes, and the deposition of these aggregates into senile plaques. The model suggests that efficient degradation of Aβ monomers alone may suffice to prevent the growth of senile plaques, even without degrading Aβ aggregates and existing plaques. This is because the degradation of Aβ monomers interrupts the supply of reactants needed for plaque formation. The impact of Aβ monomer diffusivity is demonstrated to be small, enabling the application of the lumped capacitance approximation and the derivation of approximate analytical solutions for limiting cases with both small and large rates of Aβ aggregate deposition into plaques. It is found that the rate of plaque growth is governed by two competing processes. One is the deposition rate of free Aβ aggregates into senile plaques. If this rate is small, the plaque grows slowly. However, if the rate of deposition of Aβ aggregates into senile plaques is very large, the free Aβ aggregates are removed from the intracellular fluid by deposition into the plaques, leaving insufficient free Aβ aggregates to catalyze the production of new aggregates. This suggests that under certain conditions, Aβ plaques may offer neuroprotection and impede their own growth. Additionally, it indicates that there exists an optimal rate of deposition of free Aβ aggregates into the plaques, at which the plaques attain their maximum size.

## 1 Introduction

Alzheimer’s disease (AD) is a severe neurodegenerative disorder and the leading cause of dementia (Hardy, 2006; Maqbool et al., 2016; Hung and Fu, 2017). One characteristic hallmark of AD is the spontaneous aggregation of amyloid peptides into fibrillar aggregates, aided by an autocatalytic process known as secondary nucleation (Thacker et al., 2023). The presence of accumulating Aβ deposits (senile plaques) in the brains of AD patients is a well-established fact, regardless of whether they are causative agents of the disease or simply serve as markers for it (Hardy and Selkoe, 2002; O’Brien and Wong, 2011; Selkoe and Hardy, 2016; Young-Pearse et al., 2023). Aβ monomers are produced by β- and γ-secretase cleavage of amyloid precursor protein (APP), which is synthesized in neurons (O’Brien and Wong, 2011; Hampel et al., 2021). The majority of the Aβ monomers produced are released into the extracellular environment, where they have the potential to aggregate (Rahman and Lendel, 2021).

Numerous studies have provided evidence of a link between the spread of amyloidogenic proteins and neurodegenerative diseases, including AD, Parkinson’s disease, Huntington’s disease, and Amyotrophic Lateral Sclerosis (Jucker and Walker, 2013; Klimova et al., 2015). In a study conducted by Morris et al. (2008), the Finke-Watzky (F-W) two-step model was utilized to curve-fit published data on amyloid-beta (Aβ) aggregation, obtained from studies by Bieschke et al. (2005), Vestergaard et al. (2005). Typically, the data are presented for two isoforms, Aβ40 and Aβ42, with the latter being more prone to aggregation.

Senile plaques accumulate over many years in patients with AD. The question is what processes control the rate of plaque growth. Potential candidates could include Aβ monomer diffusion, the kinetics of Aβ monomer conversion into free Aβ aggregates, the rate of Aβ monomer production and degradation, the rates of degradation of Aβ aggregates and plaques, and the rate of deposition of free Aβ aggregates into Aβ plaques.

Mathematical models aimed at understanding the intraneuronal mechanisms leading to the formation of amyloid-β plaques were developed in Kuznetsov and Kuznetsov (2018a,b), Torok et al. (2021). However, these earlier studies mainly focused on simulating intracellular processes and did not consider the extracellular diffusion of Aβ monomers, an important process to consider as Aβ monomers are easily diffusible. Kuznetsov (2024a,b) simulated the growth of Aβ plaques, neglecting and considering the diffusion of Aβ monomers, respectively. The current research builds upon previous models and proposes a model for the deposition of Aβ monomers into senile plaques.

## 2 Materials and models

### 2.1 Model equations

The F-W model is a simplified two-step process that describes the formation of certain aggregates. It involves two pseudo-elementary reaction steps: nucleation and autocatalysis. In the nucleation step, there is a continuous formation of new aggregates, while in the autocatalysis step, aggregates catalyze their own production (Morris et al., 2008; Iashchishyn et al., 2017). The two pseudo-elementary reaction steps are

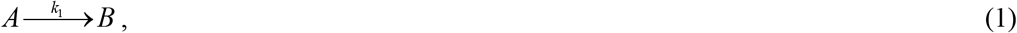

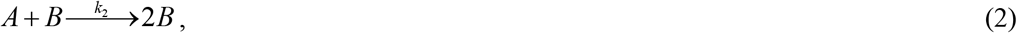

where *A* denotes a monomeric protein, and *B* denotes an amyloid-converted protein. Kinetic constants *k*^1^and *k*^2^represent the rates of nucleation and autocatalytic growth, respectively (Morris et al. (2008)). The primary nucleation process, described by Eq. (1), exclusively involves monomers. In contrast, secondary nucleation, described by Eq. (2), involves both monomers and pre-existing aggregates of the same peptide (Thacker et al., 2023).

In this study, the F-W model is employed to simulate the conversion of monomers, with a concentration denoted as *C*^*A*^, into aggregates, represented by *C*^*B*^. These aggregates encompass various forms of Aβ oligomers, protofibrils, and fibrils (Chen et al., 2017). The formation of Aβ aggregates is responsible for the deposition of amyloid plaques. Note that the minimalistic nature of the F-W model does not allow it to distinguish between different types and sizes of aggregates. Typically, the F-W model is used to investigate the conversion of a given initial concentration of monomers into aggregates. In this work, a novel extension of the F-W model is proposed to handle scenarios where monomers are continuously supplied through diffusion from the left-hand side of the system, represented by the position *x*=0 (Fig. 1). At the boundary *x*=0, which could simulate such a location as the cellular membrane (Haass et al., 2012; Masters and Selkoe, 2012), Aβ monomers are generated through the cleavage of APP (amyloid precursor protein). The boundary at *x=L* corresponds to an extracellular location. Since the entire neural tissue domain is assumed to be periodic and represented by copies of the two control volumes (CVs) shown in Fig. 1, the upper and lower boundaries of the CVs are modeled as symmetric boundaries, and there is no monomer flux in the vertical direction. Therefore, the problem is one-dimensional.

**Fig. 1.**
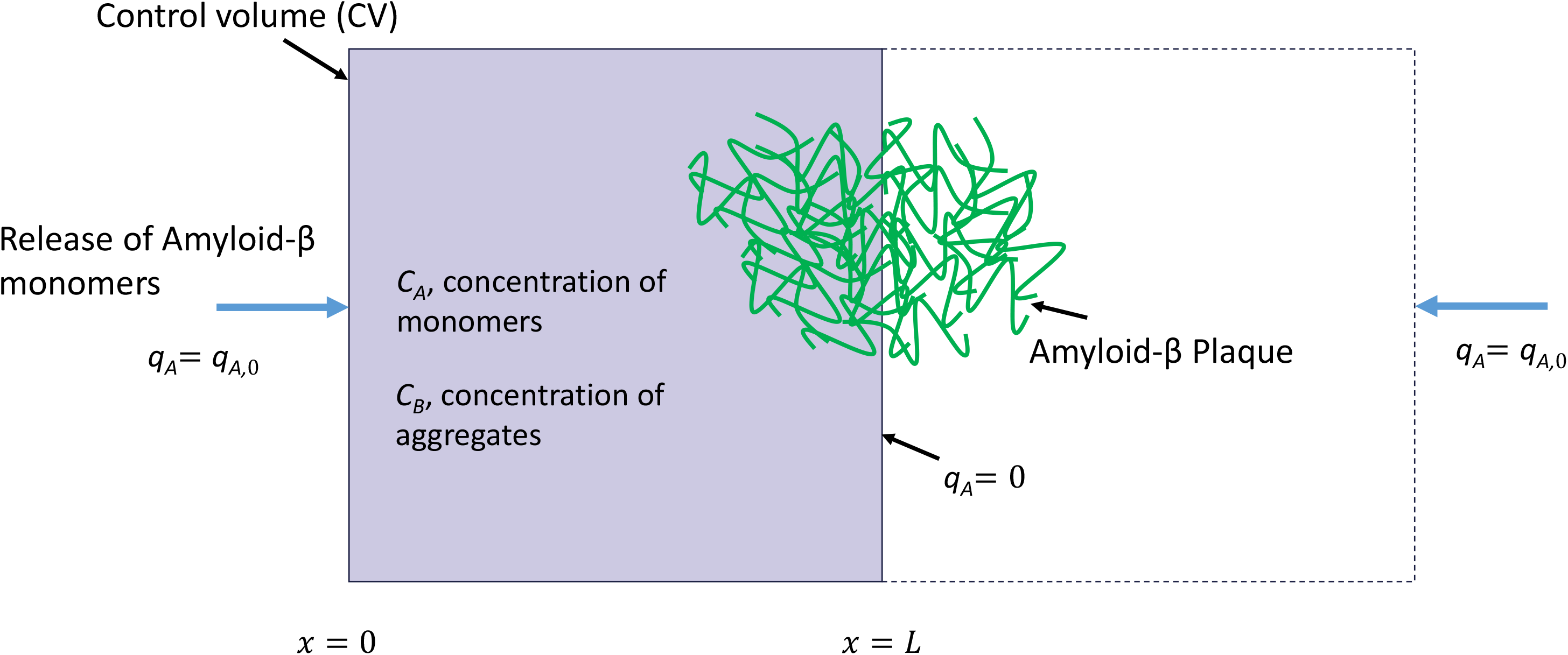
A control volume (CV) with a width of *L* = 50 μm, which is utilized to model the diffusion of Aβ monomers from the boundary at *x*=0, their accumulation as free Aβ aggregates, and subsequent deposition into a senile plaque. The boundary at *x=L* is considered to be symmetric, resulting in no flux of Aβ monomers passing through it. An Aβ plaque is assumed to form at the symmetric boundary between two CVs.

The model proposed in this paper thus is

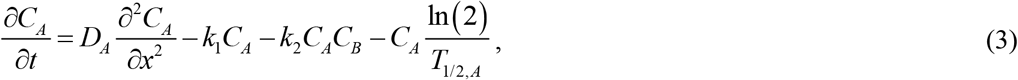

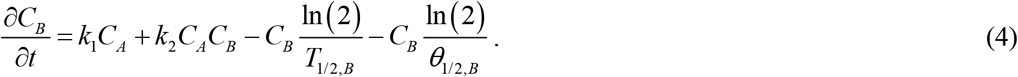

The third term on the right-hand side of Eq. (4) represents the half-life of Aβ aggregates. This term may be relevant because of the development of therapies for promoting Aβ aggregate clearance to prevent or treat AD (Yoon and Jo, 2012).

The process in which adhesive Aβ fibrils come together to form an Aβ plaque is modeled using an approach similar to that employed to model the coagulation of colloidal suspensions (Boltachev and Ivanov, 2020). It is assumed that free Aβ aggregates (*B*) deposit into Aβ plaques. The concentration of deposited aggregates is referred to as *C*_*D*_.

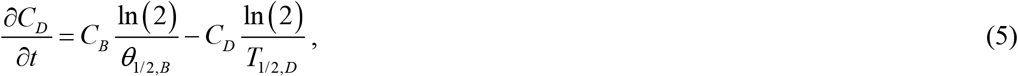

where *θ*_1/ 2,*B*_ is the half-deposition time of Aβ aggregates into Aβ plaques. This is the time it takes for half of the Aβ aggregates in the CV to be deposited into an Aβ plaque. The second term on the right-hand side of Eq. (5) represents the half-life of Aβ aggregates deposited into an Aβ plaque. This term is included due to the development of novel therapeutic interventions capable of clearing Aβ plaques (DeMattos et al., 2012).

The initial conditions are as follows:

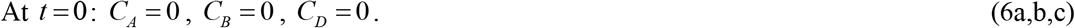

The boundary conditions are as follows:

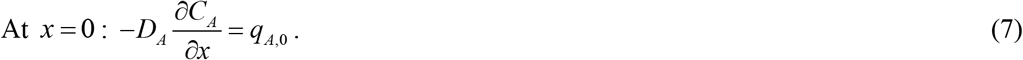

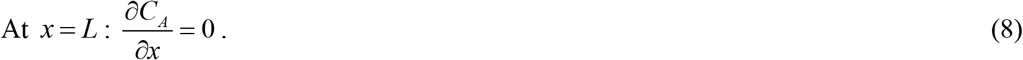

Eqs. (7) and (8) assume that Aβ monomers are released from the boundary located at *x*=0. The boundary at *x=L* is modeled as a symmetric one, as it corresponds to a half distance between senile plaques. Therefore, no flux of Aβ monomers occurs through this boundary.

The independent variables used in the model are listed in Table 1, and the dependent variables are summarized in Table 2. Table 3 provides a summary of the parameters utilized in the model.

**Table 1.** Independent variables used in the model.

| Symbol | Definition | Units |
| --- | --- | --- |
| $t$ | Time | s |
| $x$ | Linear coordinate measuring the distance from the face of the CV through which Aβ monomers are supplied | $\mu\text{m}$ |

**Table 2.** Dependent variables used in the model.

| Symbol | Definition | Units |
| --- | --- | --- |
| $C_A$ | Molar concentration of Aβ monomers | $\mu\text{M}$ |
| $C_B$ | Molar concentration of free Aβ aggregates | $\mu\text{M}$ |
| $C_D$ | Molar concentration of Aβ aggregates deposited into Aβ plaques | $\mu\text{M}$ |

**Table 3.**
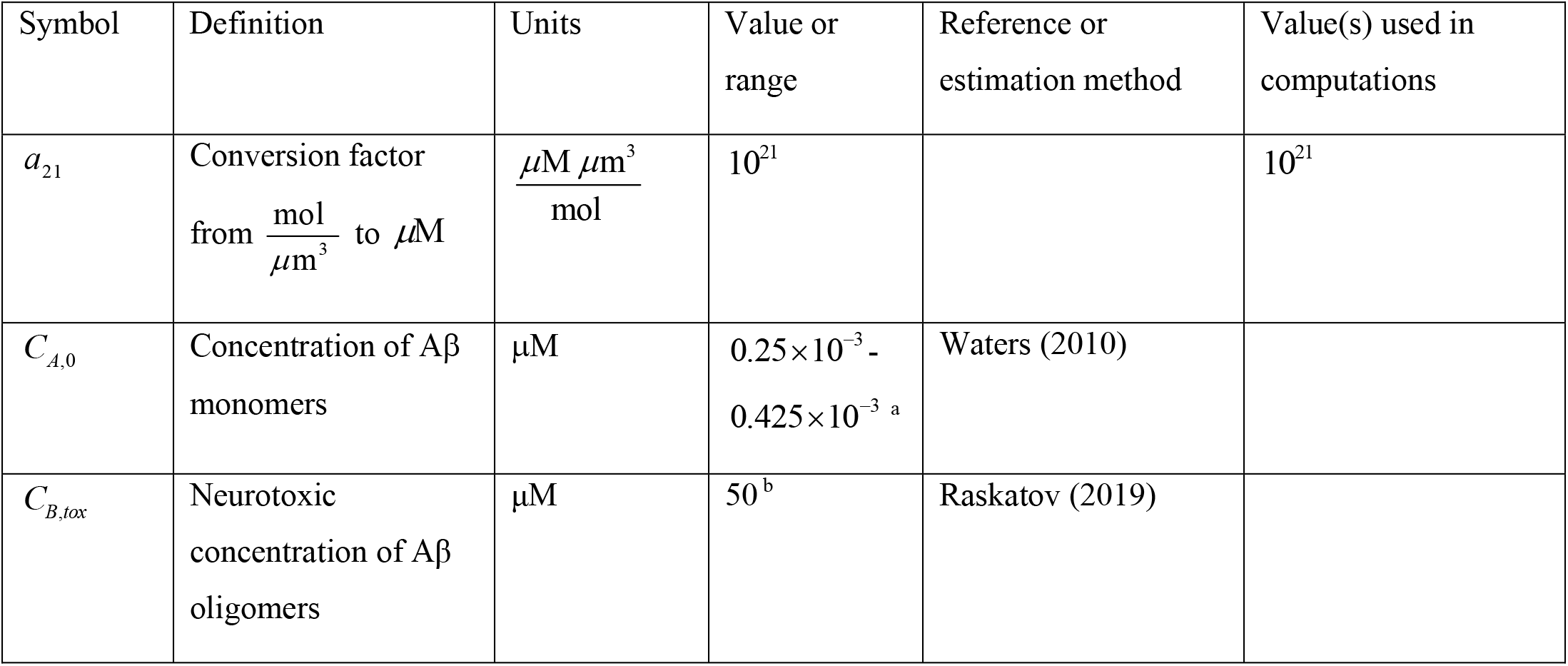
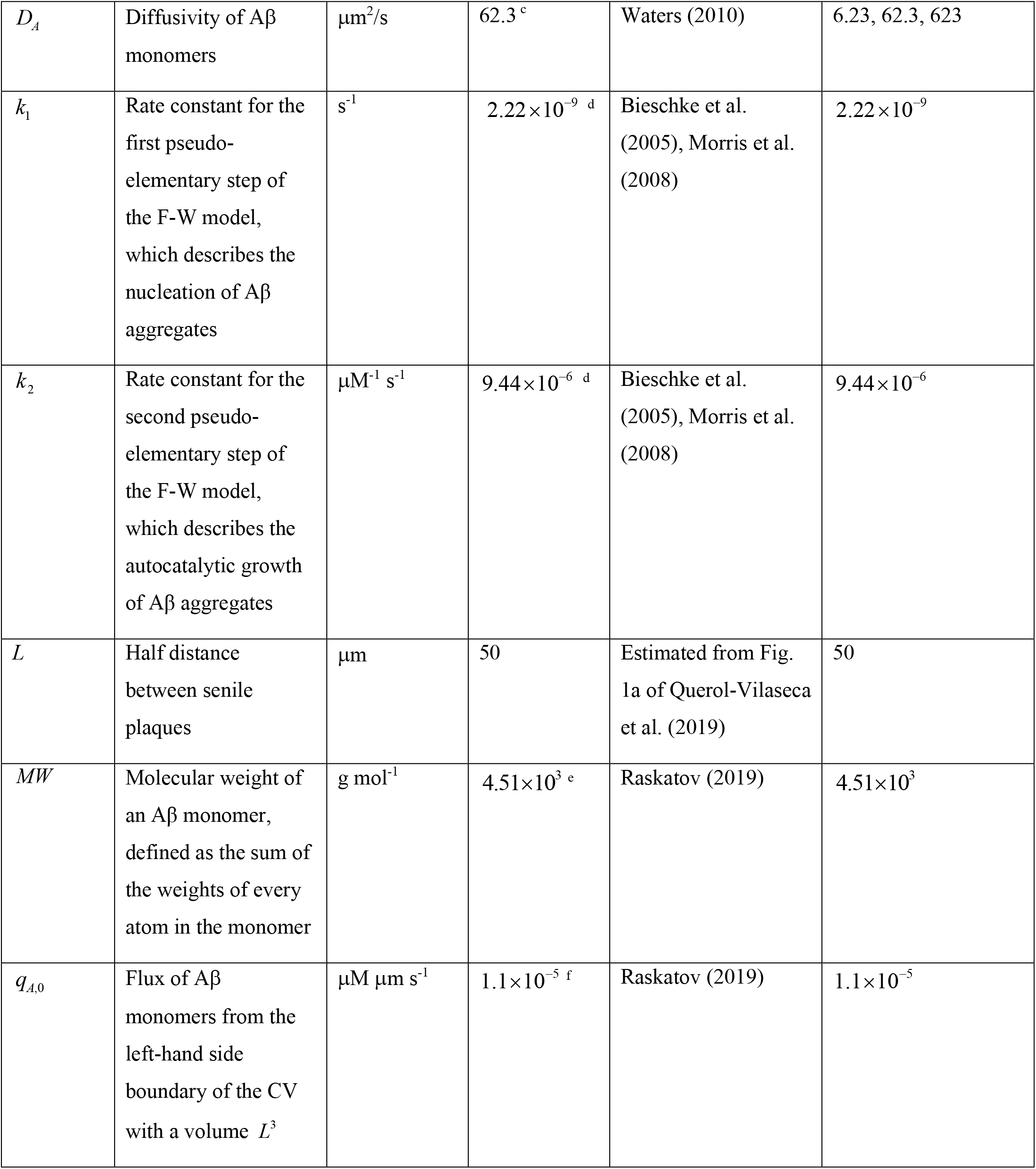
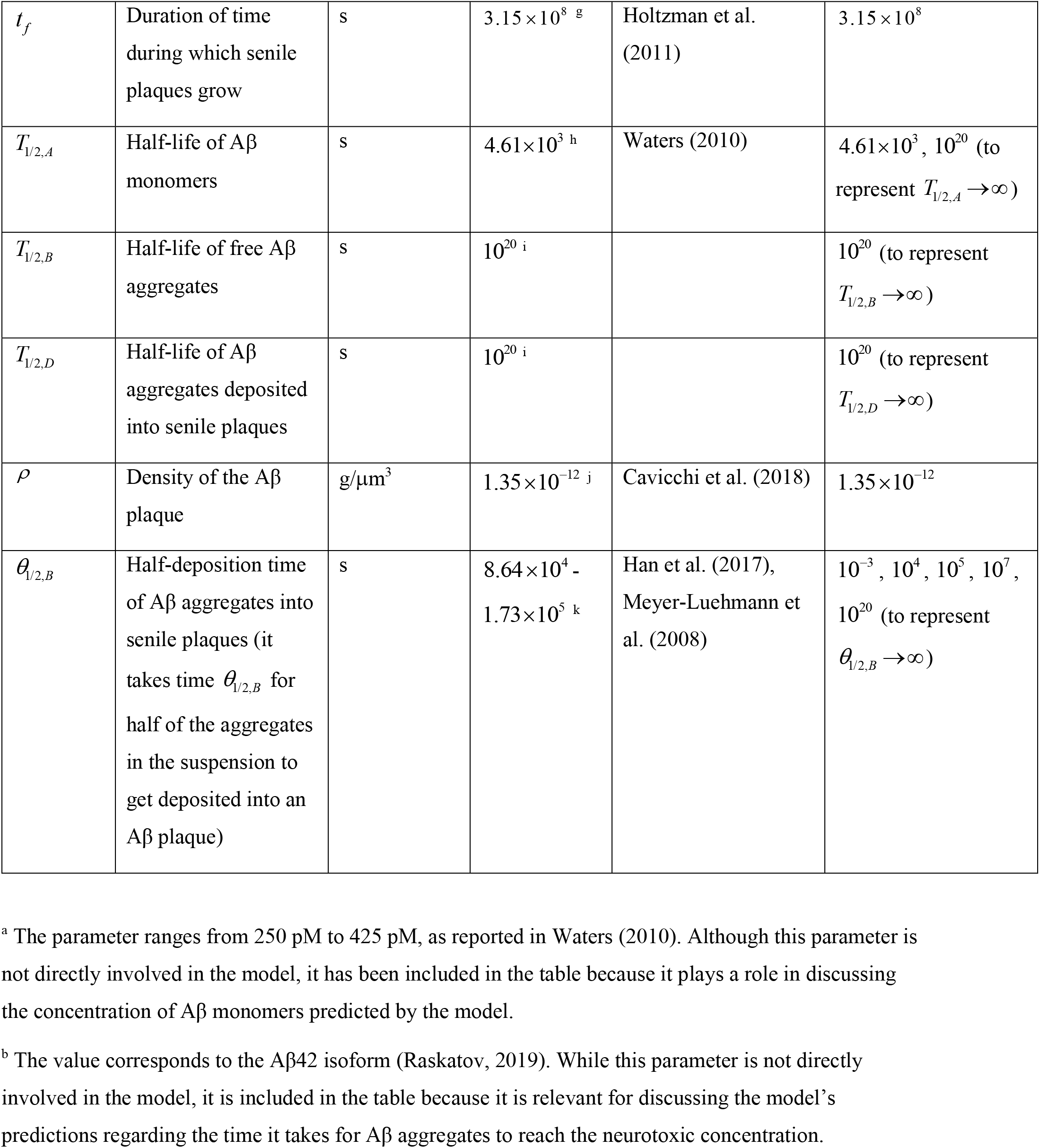
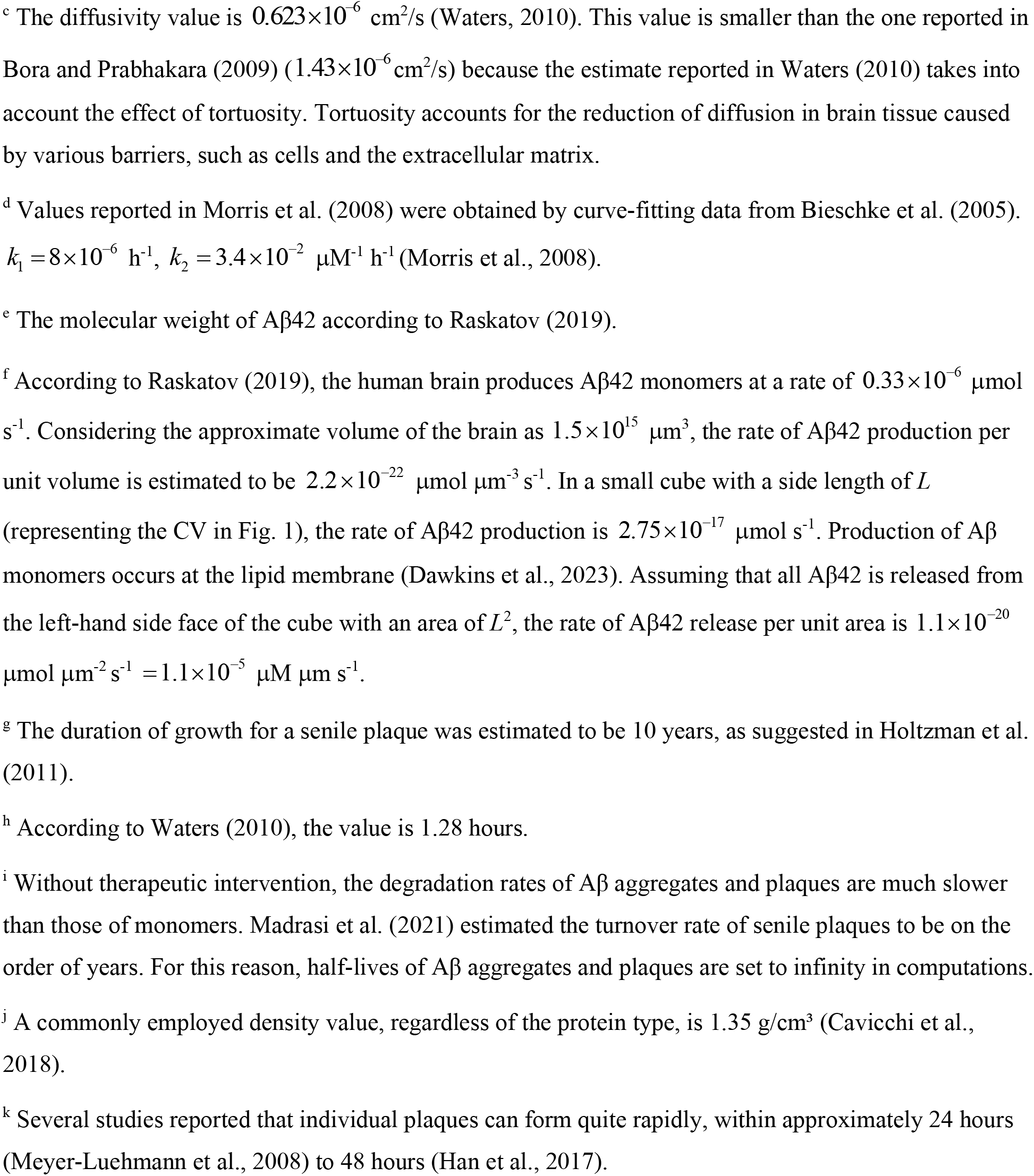
Parameters used in the model and the discussion of its results.

| Symbol | Definition | Units | Value or range | Reference or estimation method | Value(s) used in computations |
| --- | --- | --- | --- | --- | --- |
| $a_{21}$ | Conversion factor from $\frac{\text{mol}}{\mu\text{m}^3}$ to $\mu\text{M}$ | $\frac{\mu\text{M } \mu\text{m}^3}{\text{mol}}$ | $10^{21}$ | | $10^{21}$ |
| $C_{A,0}$ | Concentration of Aβ monomers | $\mu\text{M}$ | $0.25 \times 10^{-3}$ - $0.425 \times 10^{-3}$ <sup>a</sup> | Waters (2010) | |
| $C_{B,tox}$ | Neurotoxic concentration of Aβ oligomers | $\mu\text{M}$ | $50$ <sup>b</sup> | Raskatov (2019) | |
| $D_A$ | Diffusivity of Aβ monomers | $\mu\text{m}^2/\text{s}$ | 62.3 <sup>c</sup> | Waters (2010) | 6.23, 62.3, 623 |
| $k_1$ | Rate constant for the first pseudo-elementary step of the F-W model, which describes the nucleation of Aβ aggregates | $\text{s}^{-1}$ | $2.22 \times 10^{-9}$ <sup>d</sup> | Bieschke et al. (2005), Morris et al. (2008) | $2.22 \times 10^{-9}$ |
| $k_2$ | Rate constant for the second pseudo-elementary step of the F-W model, which describes the autocatalytic growth of Aβ aggregates | $\mu\text{M}^{-1} \text{s}^{-1}$ | $9.44 \times 10^{-6}$ <sup>d</sup> | Bieschke et al. (2005), Morris et al. (2008) | $9.44 \times 10^{-6}$ |
| $L$ | Half distance between senile plaques | $\mu\text{m}$ | 50 | Estimated from Fig. 1a of Querol-Vilaseca et al. (2019) | 50 |
| $MW$ | Molecular weight of an Aβ monomer, defined as the sum of the weights of every atom in the monomer | $\text{g mol}^{-1}$ | $4.51 \times 10^3$ <sup>e</sup> | Raskatov (2019) | $4.51 \times 10^3$ |
| $q_{A,0}$ | Flux of Aβ monomers from the left-hand side boundary of the CV with a volume $L^3$ | $\mu\text{M } \mu\text{m s}^{-1}$ | $1.1 \times 10^{-5}$ <sup>f</sup> | Raskatov (2019) | $1.1 \times 10^{-5}$ |
| $t_f$ | Duration of time during which senile plaques grow | s | $3.15 \times 10^8$ <sup>g</sup> | Holtzman et al. (2011) | $3.15 \times 10^8$ |
| $T_{1/2,A}$ | Half-life of Aβ monomers | s | $4.61 \times 10^3$ <sup>h</sup> | Waters (2010) | $4.61 \times 10^3$, $10^{20}$ (to represent $T_{1/2,A} \rightarrow \infty$) |
| $T_{1/2,B}$ | Half-life of free Aβ aggregates | s | $10^{20}$ <sup>i</sup> | | $10^{20}$ (to represent $T_{1/2,B} \rightarrow \infty$) |
| $T_{1/2,D}$ | Half-life of Aβ aggregates deposited into senile plaques | s | $10^{20}$ <sup>i</sup> | | $10^{20}$ (to represent $T_{1/2,D} \rightarrow \infty$) |
| $\rho$ | Density of the Aβ plaque | g/$\mu\text{m}^3$ | $1.35 \times 10^{-12}$ <sup>j</sup> | Cavicchi et al. (2018) | $1.35 \times 10^{-12}$ |
| $\theta_{1/2,B}$ | Half-deposition time of Aβ aggregates into senile plaques (it takes time $\theta_{1/2,B}$ for half of the aggregates in the suspension to get deposited into an Aβ plaque) | s | $8.64 \times 10^4$ - $1.73 \times 10^5$ <sup>k</sup> | Han et al. (2017), Meyer-Luehmann et al. (2008) | $10^{-3}$, $10^4$, $10^5$, $10^7$, $10^{20}$ (to represent $\theta_{1/2,B} \rightarrow \infty$) |

<sup>a</sup> The parameter ranges from 250 pM to 425 pM, as reported in Waters (2010). Although this parameter is not directly involved in the model, it has been included in the table because it plays a role in discussing the concentration of Aβ monomers predicted by the model.
<sup>b</sup> The value corresponds to the Aβ42 isoform (Raskatov, 2019). While this parameter is not directly involved in the model, it is included in the table because it is relevant for discussing the model's predictions regarding the time it takes for Aβ aggregates to reach the neurotoxic concentration.
<sup>c</sup> The diffusivity value is $0.623 \times 10^{-6} \text{ cm}^2/\text{s}$ (Waters, 2010). This value is smaller than the one reported in Bora and Prabhakara (2009) ( $1.43 \times 10^{-6} \text{ cm}^2/\text{s}$ ) because the estimate reported in Waters (2010) takes into account the effect of tortuosity. Tortuosity accounts for the reduction of diffusion in brain tissue caused by various barriers, such as cells and the extracellular matrix.
<sup>d</sup> Values reported in Morris et al. (2008) were obtained by curve-fitting data from Bieschke et al. (2005). $k_1 = 8 \times 10^{-6} \text{ h}^{-1}$ , $k_2 = 3.4 \times 10^{-2} \text{ } \mu\text{M}^{-1} \text{ h}^{-1}$ (Morris et al., 2008).
<sup>e</sup> The molecular weight of Aβ42 according to Raskatov (2019).
<sup>f</sup> According to Raskatov (2019), the human brain produces Aβ42 monomers at a rate of $0.33 \times 10^{-6} \text{ } \mu\text{mol s}^{-1}$ . Considering the approximate volume of the brain as $1.5 \times 10^{15} \text{ } \mu\text{m}^3$ , the rate of Aβ42 production per unit volume is estimated to be $2.2 \times 10^{-22} \text{ } \mu\text{mol } \mu\text{m}^{-3} \text{ s}^{-1}$ . In a small cube with a side length of $L$ (representing the CV in Fig. 1), the rate of Aβ42 production is $2.75 \times 10^{-17} \text{ } \mu\text{mol s}^{-1}$ . Production of Aβ monomers occurs at the lipid membrane (Dawkins et al., 2023). Assuming that all Aβ42 is released from the left-hand side face of the cube with an area of $L^2$ , the rate of Aβ42 release per unit area is $1.1 \times 10^{-20} \text{ } \mu\text{mol } \mu\text{m}^{-2} \text{ s}^{-1} = 1.1 \times 10^{-5} \text{ } \mu\text{M } \mu\text{m s}^{-1}$ .
<sup>g</sup> The duration of growth for a senile plaque was estimated to be 10 years, as suggested in Holtzman et al. (2011).
<sup>h</sup> According to Waters (2010), the value is 1.28 hours.
<sup>i</sup> Without therapeutic intervention, the degradation rates of Aβ aggregates and plaques are much slower than those of monomers. Madras et al. (2021) estimated the turnover rate of senile plaques to be on the order of years. For this reason, half-lives of Aβ aggregates and plaques are set to infinity in computations.
<sup>j</sup> A commonly employed density value, regardless of the protein type, is $1.35 \text{ g/cm}^3$ (Cavicchi et al., 2018).
<sup>k</sup> Several studies reported that individual plaques can form quite rapidly, within approximately 24 hours (Meyer-Luehmann et al., 2008) to 48 hours (Han et al., 2017).

The dimensionless form of Eqs. (3)-(8) is given in section S1.1 in Supplementary Materials. Aβ aggregation process is governed by three dimensionless parameters: (1) a parameter that defines the diffusivity of Aβ monomers, 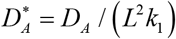; (2) a parameter that characterizes the rate at which Aβ monomers are produced, 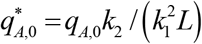; (3) a parameter that characterizes the half-life of Aβ monomers, 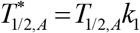; (4) a parameter that characterizes the half-life of free Aβ aggregates, 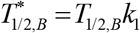; (5) a parameter that characterizes the half-life of Aβ aggregates deposited into senile plaques, 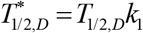; (6) a parameter that characterizes the half-deposition time of Aβ aggregates into senile plaques, 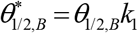.

### 2.2 Lumped capacitance approximation

#### 2.2.1 General case

A negligible variation in *C*^*A*^along *x* implies that the diffusion of Aβ monomers is infinitely fast. One of the supporting arguments for this assumption is that if the soluble Aβ diffuses slower than the speed of aggregation, growth will happen at the aggregate’s tips, resulting in a ramified tree-like structure, which contrasts with the experimental observations (Cruz et al., 1997). Under this assumption, the CV shown in Fig. 1 can be treated as a lumped capacitance body and Eqs. (3)-(5) can be reformulated as follows:

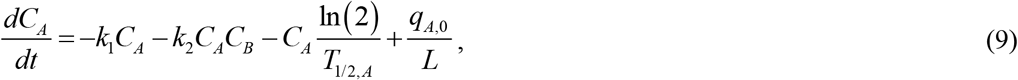

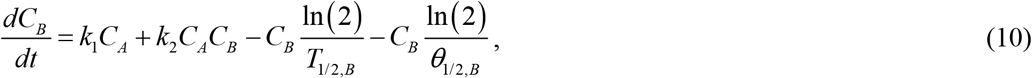

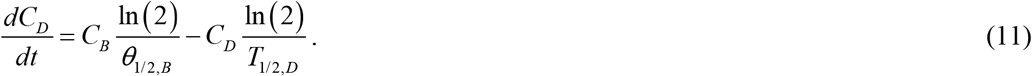

Kuznetsov (2024b) proposed the following dimensionless parameter

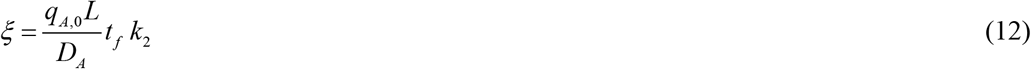

and demonstrated that the lumped capacitance approximation is valid when *ξ* ≪ 1. In Eq. (12) *t* _*f*_ represents the process duration.

The dimensionless forms of Eqs. (9)-(11) are given in section S1.2 in Supplementary Materials.

#### 2.2.2 The case when Aβ aggregates quickly deposit into senile plaques, *θ*_1/ 2,*B*_→ 0

For *θ*_1/ 2,*B*_→ 0, which physically means immediate deposition of Aβ aggregates into plaques, Eqs. (9)-(11) collapse to

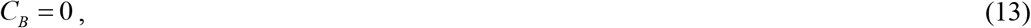

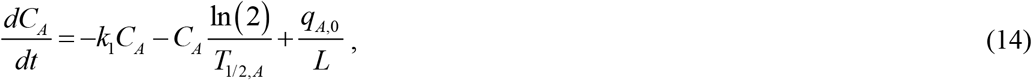

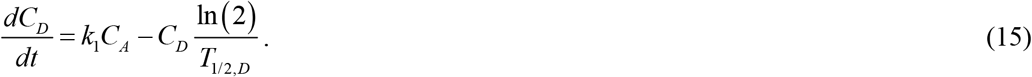

Upon adding Eqs. (14) and (15) for the case of *T*_1/2, *A*_→∞ and *T*_1/2,*D*_→∞, the following result is obtained:

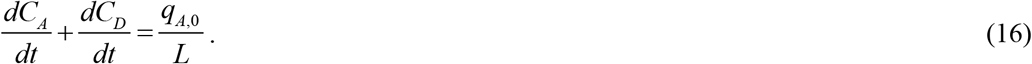

Integrating Eq. (16) with respect to time and using the initial condition given by Eq. (6a,c), the following result is obtained:

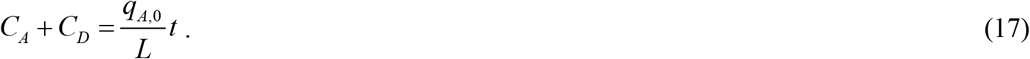

The increase of *C*_*A*_+ *C*_*D*_ with time is attributed to the supply of Aβ monomers through the left-hand side boundary of the CV. By eliminating *C*_*A*_ from Eq. (15) using Eq. (17), the following is obtained:

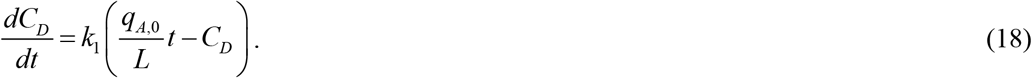

The exact solution of Eq. (18) with the initial condition given by Eq. (6c) is as follows:

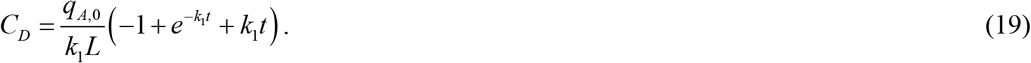

If *t* → 0, Eq. (19) implies that

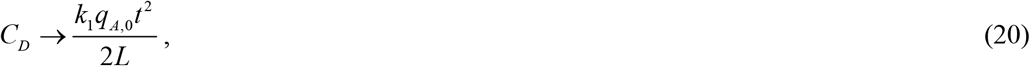

indicating that at small *t*, the concentration of Aβ aggregates deposited into Aβ plaques is directly proportional to the kinetic constant that describes the nucleation of Aβ aggregates.

*C*_*A*_ can then be found using Eqs. (17) and (19) as:

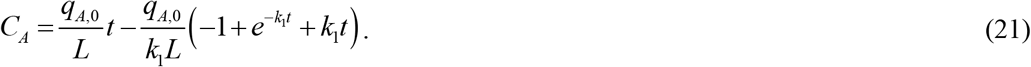

#### 2.2.3 The case of a slow rate of deposition of Aβ aggregates into senile plaques, *θ*_1/2,*B*_→∞

For *θ*_1/2,*B*_→∞, which physically implies that Aβ aggregates do not deposit into senile plaques at all, Eqs. (9)-(11) collapse to

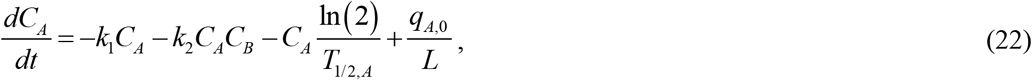

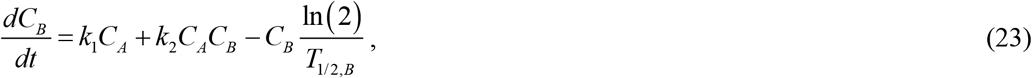

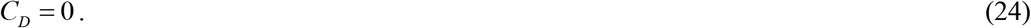

Eqs. (22) and (23) were solved with the initial conditions (6a,b) in dimensionless form, as described in Kuznetsov (2024a). To ensure that the paper is self-contained, the analytical solution of Eqs. (22) and (23) is provided in section S1.2 of the Supplementary Materials.

#### 2.2.4 Developing a model for senile plaque growth

The method employed to simulate the growth of an Aβ plaque (see Fig. 1) involves calculating the total number of Aβ monomers, denoted as *N*, integrated into the plaque at a given time *t*. This calculation is based on the equation proposed in Watzky et al. (2008):

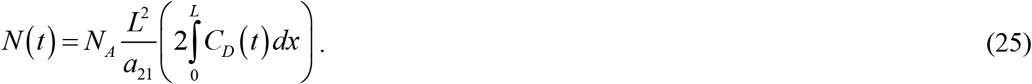

The factor of 2 preceding the integral in Eq. (25) is included based on the assumption of the model that the Aβ plaque develops between two neurons, with each membrane releasing Aβ monomers at a rate of *q*_*A*,0_(Fig. 1).

Alternatively, *N* (*t*) can be found according to Watzky et al. (2008) as follows:

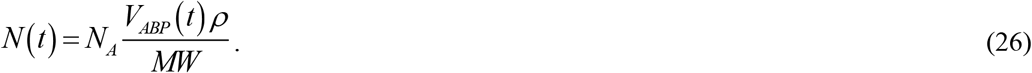

Here, *MW* represents the mean molecular weight of an Aβ monomer.

By equating the expressions on the right-hand sides of Eqs. (25) and (26) and subsequently solving for the volume of an Aβ plaque, the following result is obtained:

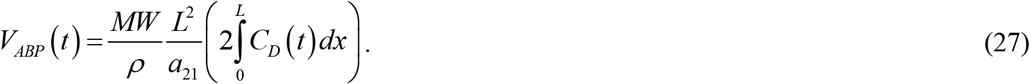

Here, 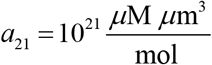 represents the conversion factor from 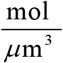 to *μ*M.

Assuming the Aβ plaque to be spherical,

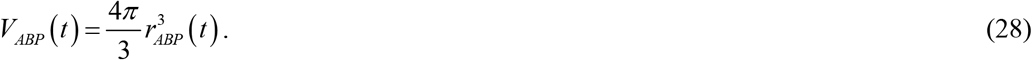

By equating the expressions on the right-hand sides of Eqs. (27) and (28) and solving for the radius of the Aβ plaque, the following result is obtained:

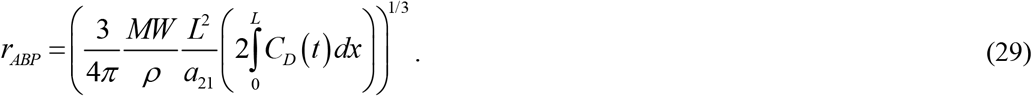

When employing the lumped capacitance approximation and substituting the approximate solution for *C*_*D*_ from Eq. (19) into Eq. (29), the following approximate solution for *r*_*ABP*_ is derived:

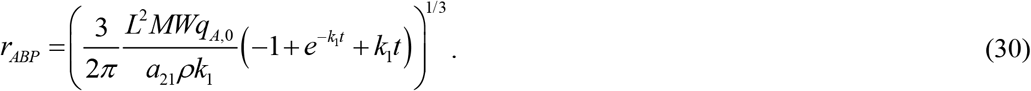

If *t* → 0, Eq. (30) implies that

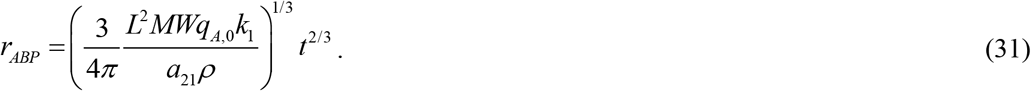

### 2.3 Sensitivity analysis

One of the main objectives of this study was to investigate the sensitivity of the radius of an Aβ plaque to several factors, including the half-deposition time of Aβ aggregates into senile plaques, *θ*_1/2,*B*_ the diffusivity of Aβ monomers, *D*_*A*_; the rate of Aβ monomer production, characterized by *q*_*A*,0_; the rate constant describing the nucleation of Aβ aggregates, *k*_1_; and the rate constant describing the autocatalytic growth of Aβ aggregates, *k*_2_.

This involved calculating the local sensitivity coefficients, which represent the first-order partial derivatives of the observable (the radius of the Aβ plaque) with respect to parameters such as the half-deposition time of Aβ aggregates into senile plaques. This calculation was conducted following the approach developed in Beck and Arnold (1977), Zadeh and Montas (2010), Zi (2011), Kuznetsov and Kuznetsov (2019). Specifically, the sensitivity coefficient of *r*_*ABP*_to (for example) *θ*_1/2,*B*_was determined as follows:

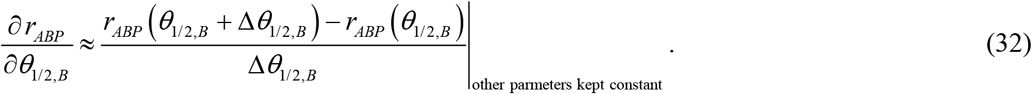

Here, Δ*θ*_1/ 2,*B*_= 10^−3^*θ*_1/ 2, *B*_ is the step size. Calculations were performed using different step sizes to assess the sensitivity coefficients’ independence from the step size.

The non-dimensionalized relative sensitivity coefficients were computed according to the method outlined by Zadeh and Montas (2010), Kacser et al. (1995), as, for example:

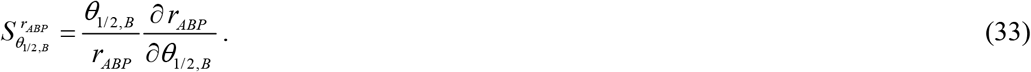

From Eq. (30), it follows that

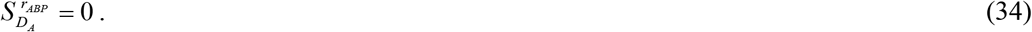

Eq. (34) is expected because Eq. (30) is obtained under the lumped capacitance approximation, which assumes that the diffusivity of Aβ is infinitely large. Similarly, using Eq. (30), the following is obtained:

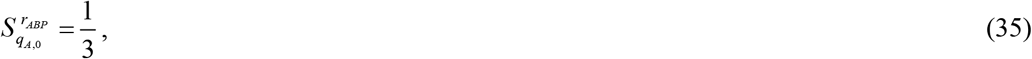

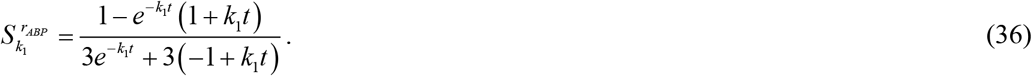

If *k*_1_→ 0, Eq. (36) leads to

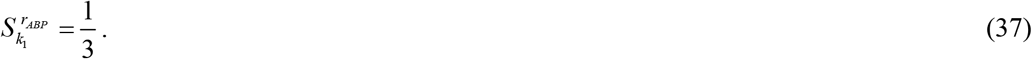

Furthermore,

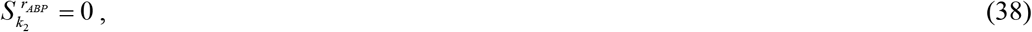

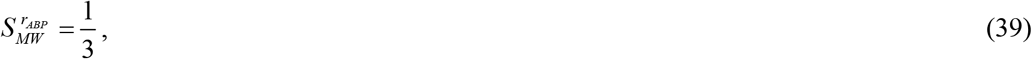

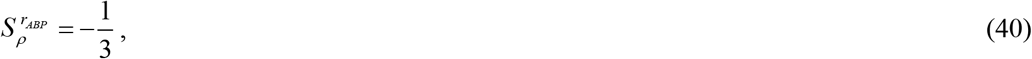

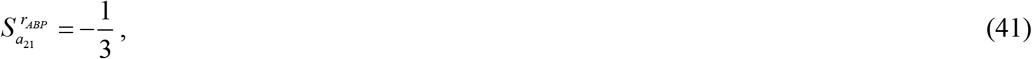

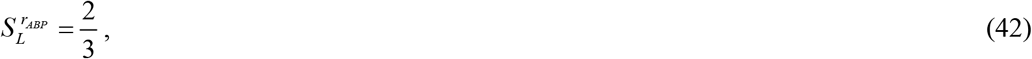

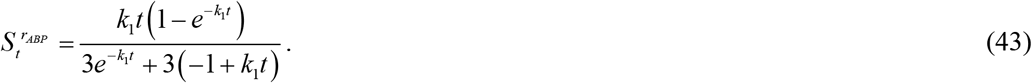

If *t* → 0, Eq. (43) gives that

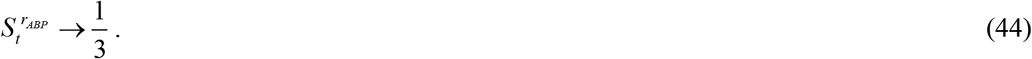

## Results

Details of the numerical solution are given in section S1.3 of the Supplementary Materials. Fig. 2 depicts the scenario where the half-life of Aβ monomers is assumed to be infinite. The concentrations of monomers, *C*_*A*_, free aggregates, *C*_*B*_, and aggregates deposited into Aβ plaques, *C*_*D*_, gradually decrease from *x*=0 to *x=L*. This decline occurs due to monomer diffusion from the boundary where monomers are supplied (*x*=0), leading to the highest concentration, to the boundary with a symmetric (zero flux) boundary condition (*x=L*), where Aβ monomer and both types of aggregate concentrations are at their lowest (Fig. 2). Note that in this case, the concentration of aggregates deposited into Aβ plaques reaches 19.5 μM within 10 years, which is in the same order of magnitude as the neurotoxic level of 50 μM (Table 3). The scenario where the half-life of Aβ monomers is finite, *T*^1/ 2, *A*^ = 4.61×10^3^ s, is displayed in Fig. S1. In Fig. S1a, the concentration of Aβ monomers, *C*_*A*_, is approximately 1.465×10^−3^ μM, which is of the same order of magnitude as the range reported in Waters (2010) (Table 3): 0.25×10^−3^ to 0.425×10^−3^ μM. Note that the concentration of Aβ aggregates deposited into Aβ plaques, *C*_*D*_, is approximately 1.025×10^−3^ μM (Fig. S1), which is four orders of magnitude less than the neurotoxic concentration of 50 μM (Table 3). This suggests that neurotoxicity of Aβ and the formation of Aβ plaques are expected to occur only when the machinery responsible for Aβ monomer degradation is not functioning properly. This suggests that AD therapies should focus not only on the clearance of Aβ aggregates but also of Aβ monomers because, unless cleared, they may assemble into aggregates (Zuroff et al., 2017).

**Fig. 2.**
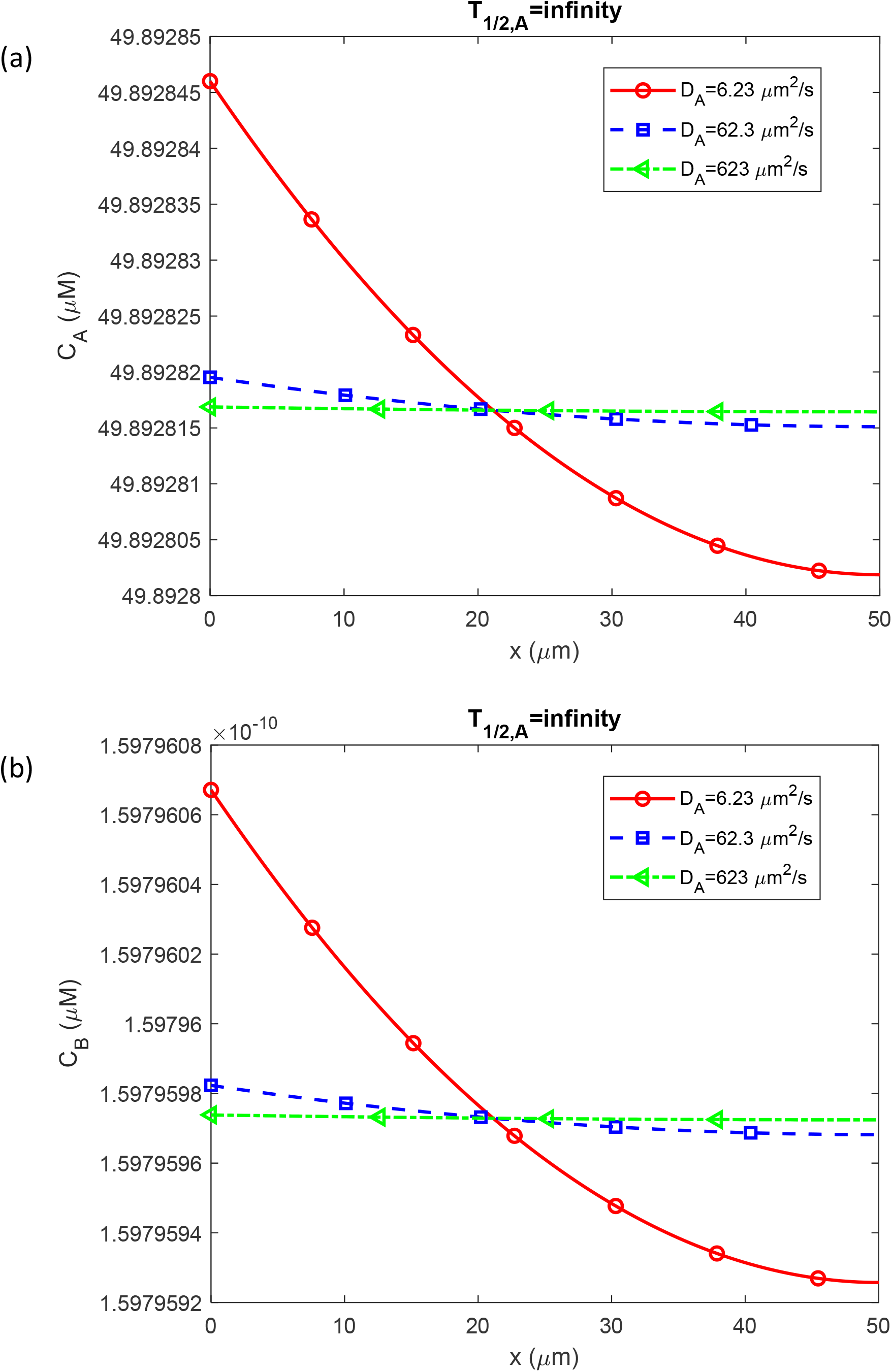
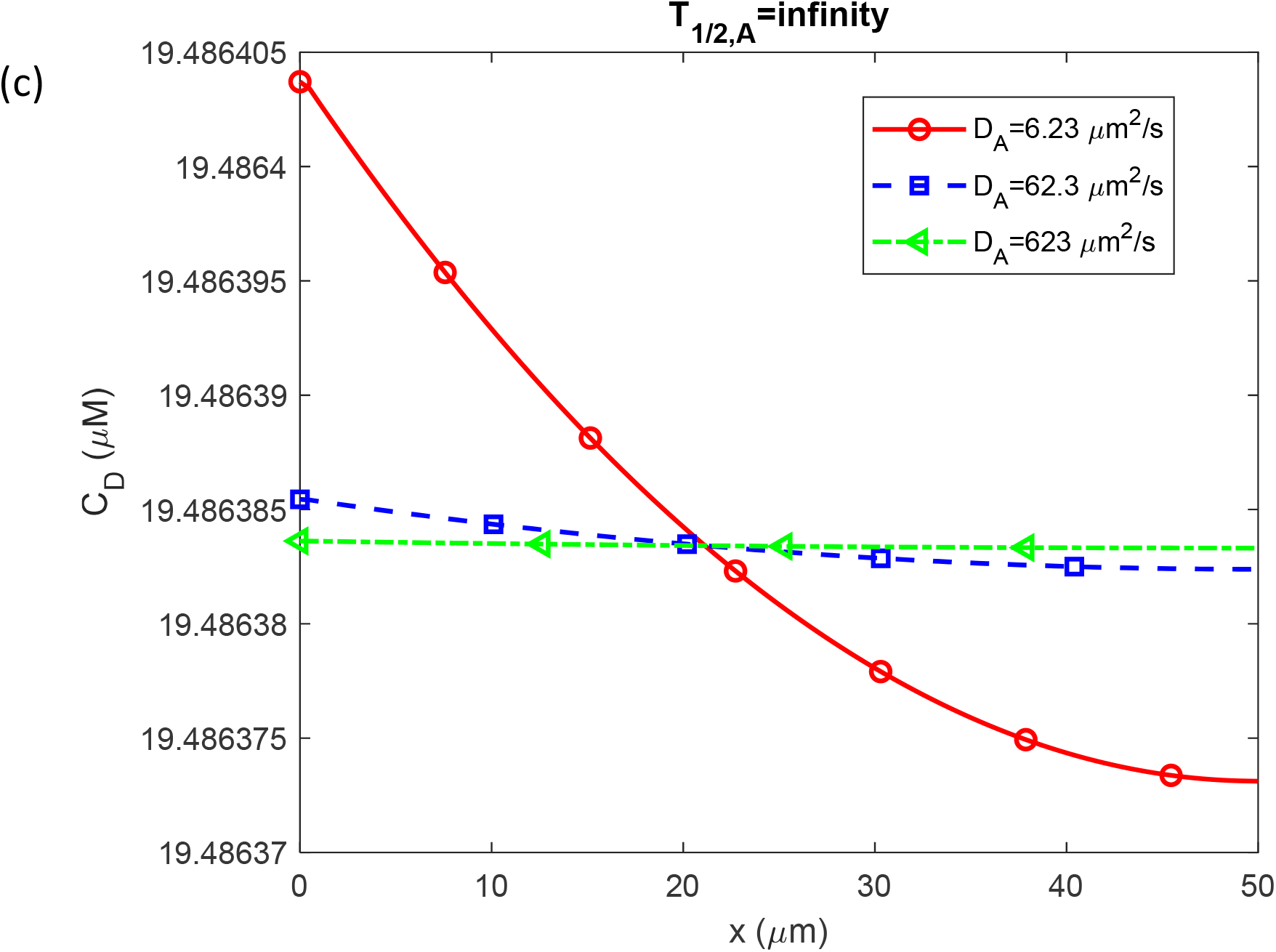
Molar concentration of Aβ monomers, *C*_*A*_(a); free Aβ aggregates, *C*_*B*_(b); and Aβ aggregates deposited into Aβ plaques, *C*_*D*_(c); as a function of the distance from the surface that releases Aβ monomers (e.g., cell membrane). These computational results are presented at *t* = 3.15×10^8^ s =10 yearsand correspond to the scenario with an infinite half-life of Aβ monomers and a small half-deposition time of Aβ aggregates into senile plaques, *θ*_1/2,*B*_=10^−3^ s.

Figs. 2 and S1 show that the variations of the concentrations across the CV compared to their average values do not exceed 0.0001%. Thus, the lumped capacitance approximation is applicable.

Figs. S2 and S3 depict concentrations of Aβ monomers and aggregates vs. time for the infinite and finite half-life of Aβ monomers, respectively. Initially, at the right-hand side boundary of the CV, where *x* = *L*, the concentrations of Aβ monomers and aggregates are zero, as indicated by the initial condition in Eq. (6). Subsequently, all three concentrations increase due to the continuous supply of reactants (monomers) through the left-hand side boundary of the CV at *x*=0, followed by their conversion into free aggregates and deposition of the free aggregates into amyloid plaques (Figs. 2 and 3). When the diffusivity of the monomers is decreased, the concentration of aggregates deposited into Aβ plaques exhibits a slightly slower increase, as illustrated by the red solid curve in Fig. S3c corresponding to *D*_*A*_= 6.23 μm^2^/s. Despite this very slight divergence, the curves in Figs. S2 and S3 computed for different values of the diffusivity of Aβ monomers almost coincide, which reaffirms that using the lumped capacitance approximation for this problem is reasonable. To confirm this, equations obtained under the lumped capacitance approximation were solved analytically for the case when Aβ aggregates immediately deposit into Aβ plaques, the scenario modeled by *θ*_1/2,*B*_ →0. The curves depicting approximate solutions (referred to as such due to the lumped capacitance approximation) for *C*_*A*_ and *C*_*D*_, given by Eqs. (21) and (19) respectively, coincide with the curves obtained by numerically solving Eqs. (3)-(8) (Fig. 2a,c).

**Fig. 3.**
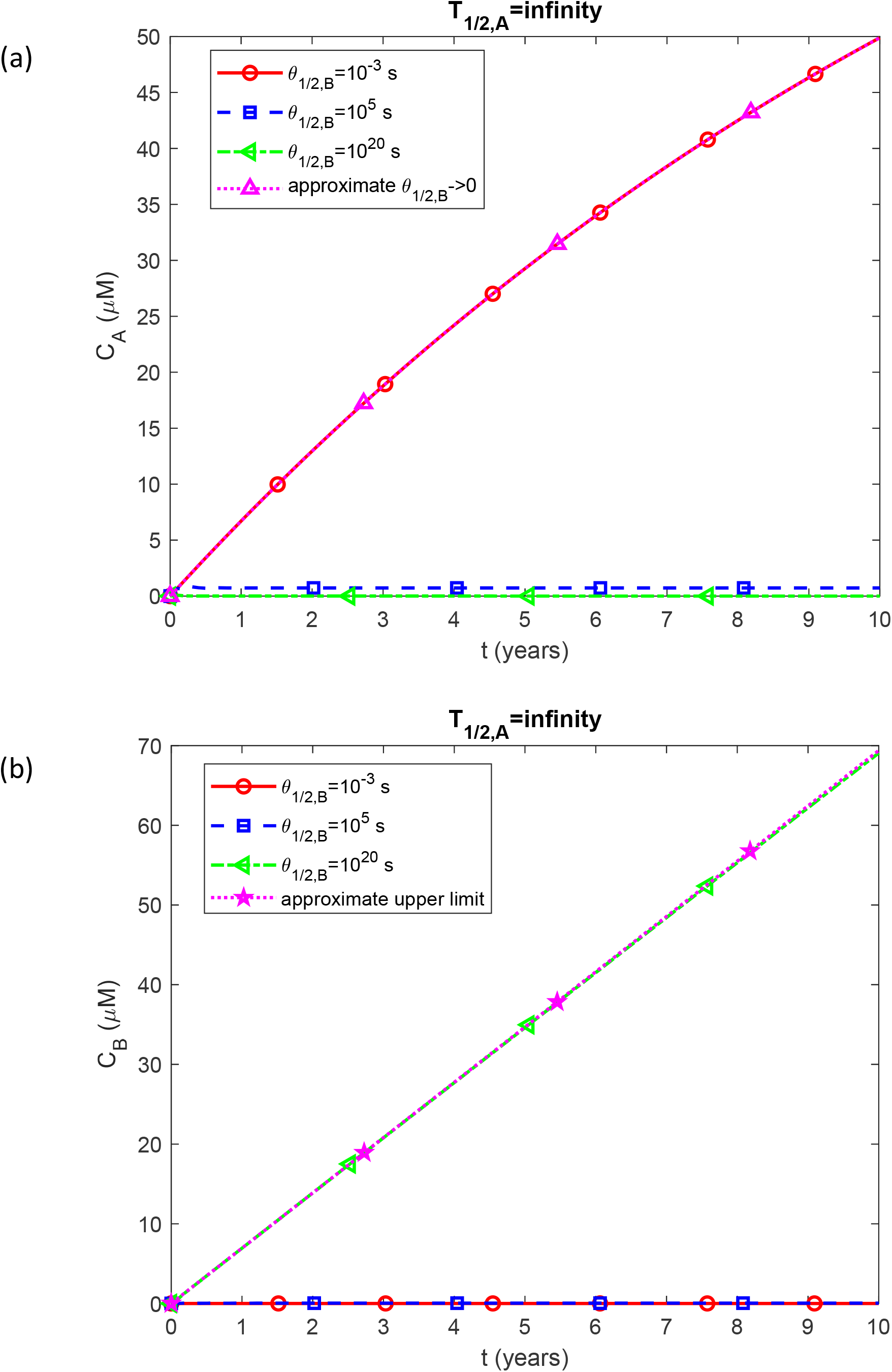
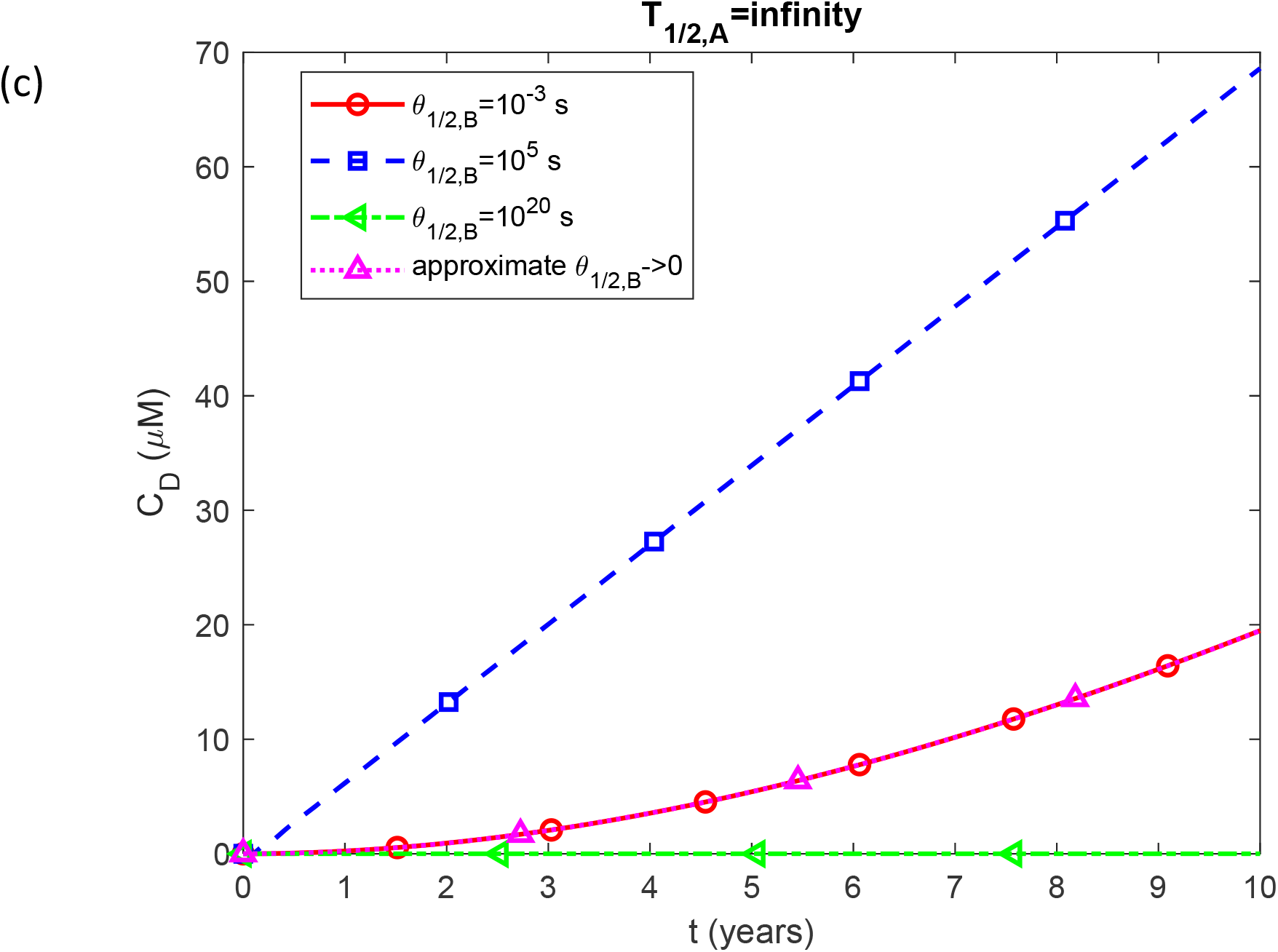
(a) Molar concentration of Aβ monomers, *C*_*A*_(a); free Aβ aggregates, *C*_*B*_(b); and Aβ aggregates deposited into Aβ plaques, *C*_*D*_(c); as a function of the half-deposition time of Aβ aggregates into senile plaques, *θ*_1/2,*B*_. The computational results are presented at the right-hand side boundary of the CV, *x=L*, at *t* = 3.15×10^8^ s =10 years. The depicted scenario assumes an infinite half-life of Aβ monomers.

If the half-life of Aβ monomers is assumed to be finite, *T*_1/2, *A*_ = 4.61×10^3^ s (Figs. S3), the concentration of Aβ aggregates at *t*=10 years decreases significantly compared to the scenario with an infinite half-life, from approximately 19 μM to approximately 10^−3^ μM (compare Fig. S3c with Fig. S2c). This notable decrease in the aggregate concentration occurs because, with a smaller half-life, a significant portion of the monomers is degraded by the proteolytic system rather than being converted into aggregates. This finding has significant implications for AD. Many studies have linked AD to the failure of the proteolytic machinery to maintain low Aβ concentrations and to degrade faulty and misfolded Aβ (Wang et al., 2006; Baranello et al., 2015; Kim and Priefer, 2020). The results obtained from this study explain why preventing Aβ aggregation requires more than just degrading faulty Aβ. A properly functioning proteolytic machinery capable of degrading Aβ monomers, which are constantly being produced, can protect the brain from Aβ aggregates. Interestingly, to achieve this protection, the ability to degrade aggregates is not required (notably, *T*_1/2,*B*_ and *T*_1/2,*D*_ were set to 10^20^ s (Table 3), which simulates the scenario with no degradation of free and deposited Aβ aggregates).

The radius of a growing Aβ plaque, *r*_*ABP*_, vs time for the infinite and finite half-life of Aβ monomers is depicted in Fig. S4 and S5, respectively. For the scenario with dysfunctional degradation machinery for Aβ monomers (*T*^1/2,*A*^→∞), the radius of the Aβ plaque predicted by Eq. (30), which is obtained with the lumped capacitance approximation, coincides with numerically obtained curves (Fig. S4). The plaque radius for *T*_1/2, *A*_→∞ reaches approximately 1.5 μm after 10 years of growth (Fig. S4). This is an order of magnitude less than the representative diameter of an Aβ plaque observed in Querol-Vilaseca et al. (2019) (50 μm). However, plaques can vary in size. Also, the plaque diameter can be easily increased in the model by using a growth time larger than 10 years (for example, 20 years), increasing the production rate of Aβ monomers, *q*_*A*,0_, and increasing the distance between the plaques, *L*. Importantly, for the scenario with normally functioning degradation machinery, *T*_1/ 2, *A*_ = 4.61×10^3^, the radius of the Aβ plaque reaches only approximately 0.06 μm (Fig. S5). This suggests that the formation of Aβ plaques of detectable size is only possible if the monomer degradation machinery is dysfunctional. In the scenario with an infinitely large half-life of Aβ monomers (Fig. 3), the concentration of monomers, *C*_*A*_, exhibits the fastest growth when free aggregates are immediately deposited into Aβ plaques (*θ*_1/2,*B*_=10^−3^), see Fig. 3a. This occurs because the conversion of Aβ monomers into aggregates is catalyzed by the aggregates themselves (see Eqs. (3) and (4)). If free aggregates are immediately deposited into plaques, the concentration of aggregates capable of catalysis is very low (as depicted by the red solid line in Fig. 3b). The monomers, however, are constantly supplied from the left side of the CV (refer to Fig. 1). This explains the rapid increase of *C*_*A*_ in Fig. 3a in the case of *θ*_1/2,*B*_ =10^−3^ s. The numerically computed curve for this scenario coincides with the approximate solution in the case of *θ*_1/2,*B*_→0 given by Eq. (21) (refer to Fig. 3a). On the other hand, if free aggregates are never deposited into plaques (the case of *θ*_1/2,*B*_ =10^20^ s), the large number of catalytically active aggregates in the intracellular fluid leads to monomers being quickly converted into aggregates. This prevents the concentration of monomers from increasing despite their continuous production, as illustrated by the green dash-dot line in Fig. 3a.

If Aβ never deposits into Aβ plaques (the case of *θ*_1/2,*B*_ =10^20^ s), the concentration of free Aβ aggregates, *C*_*B*_, linearly increases as these aggregates are continuously produced by the conversion from monomers, as shown by the green dash-dot line in Fig. 3b. The line depicting this case coincides with the line computed using the approximate solution given by Eq. (S14) (Fig. 3b).

Fig. 3c illustrates an interesting competition between two effects. If free Aβ aggregates do not deposit into plaques (*θ*_1/2,*B*_ =10^20^ s), the concentration of Aβ aggregates deposited into plaques, *C*_*D*_, is zero. If Aβ aggregates deposit into the plaques very quickly (*θ*_1/2,*B*_=10^−3^s), the concentration of Aβ aggregates deposited into the plaques, *C*_*D*_, is also quite low because free aggregates are needed to catalyze the conversion of monomers into free aggregates (see Eq. (2)). Therefore, *C*_*D*_grows at the fastest rate at an intermediate value of *θ*_1/ 2, *B*_, as indicated by the blue dashed line corresponding to *θ*_1/ 2, *B*_ =10^5^ s. Also, note that the numerical curve in the case of *θ*_1/2,*B*_=10^−3^ s coincides with that computed using Eq. (19) in the case of *θ*_1/2,*B*_→0.

The scenario with a physiologically relevant half-life of Aβ monomers, *T*_1/ 2, *A*_ = 4.61×10^3^ s, predicts much smaller concentrations because many monomers are destroyed before they have a chance to be converted into aggregates (Fig. S6). The largest concentration of free Aβ aggregates, *C*_*B*_, is in the case when free aggregates never deposit into the plaques, *θ* _1/2,*B*_=10^20^ s (Fig. S6b). The largest concentrations of Aβ aggregates deposited into plaques, *C*_*D*_, are observed in the cases when free aggregates quickly deposit into the plaques (*θ*_1/2,*B*_=10^−3^ s and *θ*_1/2,*B*_ =10^5^ s).

The radius of the Aβ plaque computed in the case of an infinitely large half-life of Aβ monomers is the largest for the intermediate value of the half-deposition time of Aβ aggregates into senile plaques, *θ*_1/2,*B*_=10^5^ s (Fig. 4), which is consistent with the results in Fig. 3c. Note that the numerical curve computed for this case coincides with the curve computed using the approximate solution given by Eq. (S15), which predicts the increase of the plaque radius proportionally to the cube root of time. Also note that the curve depicting the case of very fast deposition of Aβ aggregates into the senile plaques (*θ*_1/2,*B*_=10^−3^ s) coincides with that computed using the approximate solution given by Eq. (30).

**Fig. 4.**
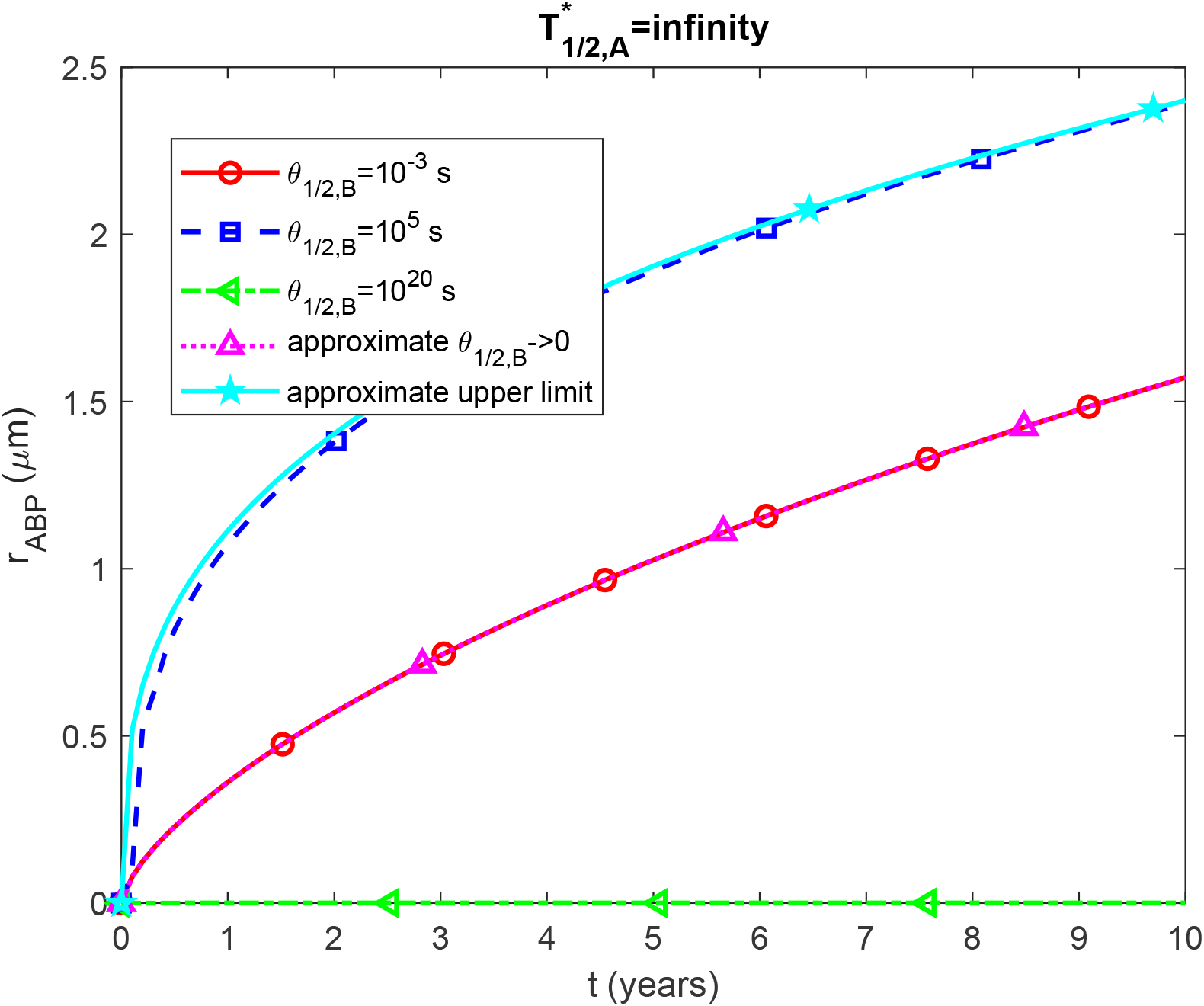
(a) Radius of a growing Aβ plaque, *r*_*ABP*_, vs time (in years). The numerical solution is obtained by numerically solving the problem given by Eqs. (3)-(8) and then using Eq. (29) to calculate the radius of the Aβ plaque. The scenario displayed in this figure assumes an infinite half-life of Aβ monomers. The curve marked as “approximate *θ*_1/2,*B*_ →0” is computed utilizing Eq. (30). The curve marked as “approximate upper limit” is computed utilizing Eq. (S15).

In the case of a physiologically relevant half-life of Aβ monomers, *T*_1/ 2, *A*_ = 4.61×10^3^ s (Fig. S7), the radius grows much slower than in the case of an infinite half-life (Fig. 4), which reaffirms the hypothesis that senile plaque growth can occur only with dysfunctional degradation machinery for Aβ monomers.

The results presented in Figs. 3 and 4 served as motivation to investigate how the Aβ monomer and aggregate concentrations depend on the half-deposition time of Aβ aggregates into senile plaques, *θ*_1/2,*B*_(Fig. 5). In the case of an infinitely large half-life of Aβ monomers, the concentration of monomers, *C*_*A*_, decreases from a value of approximately 50 μM, as predicted by Eq. (21), to a value close to 0, as predicted by Eq. (S13), within the range of *θ*_1/2,*B*_ between 10^1^ −10^5^ s. The results are practically independent of the value of *D*_*A*_(Fig. 5a).

**Fig. 5.**
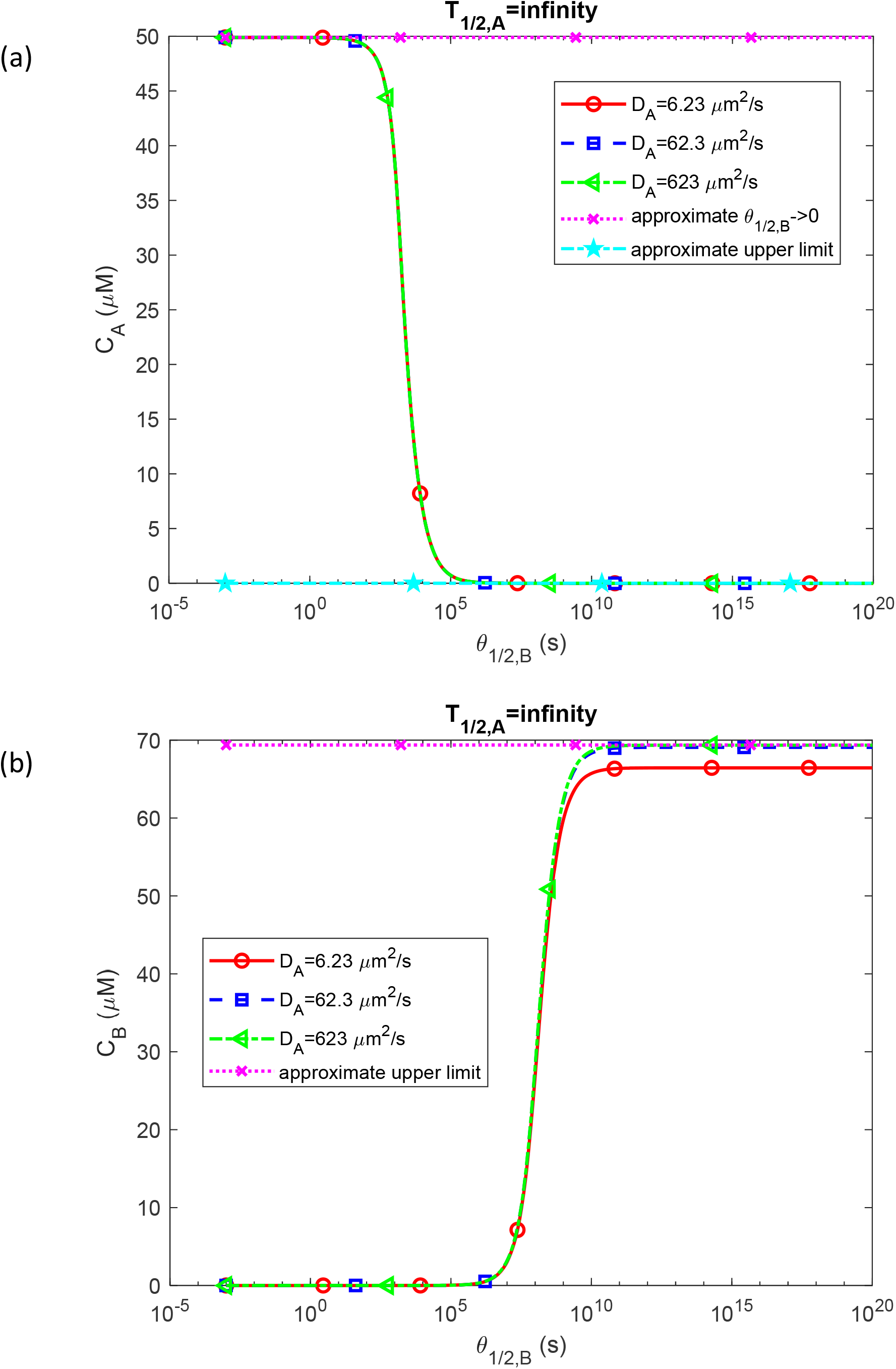
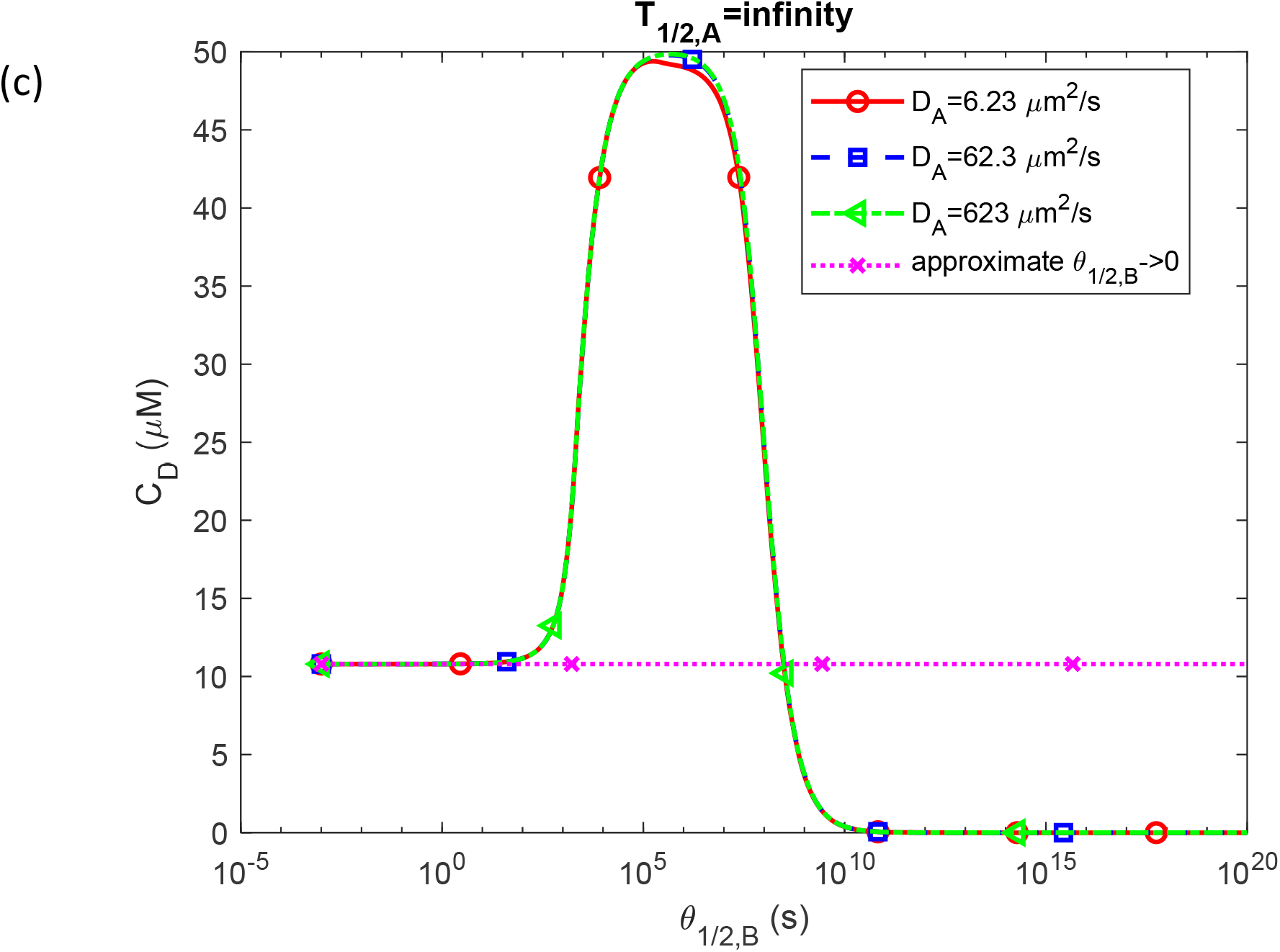
Molar concentration of Aβ monomers, *C*_*A*_(a); free Aβ aggregates, *C*_*B*_(b); and Aβ aggregates deposited into Aβ plaques, *C*_*D*_(c); as a function of the half-deposition time of Aβ aggregates into senile plaques, *θ*_1/2,*B*_. The computational results are presented at the right-hand side boundary of the CV, *x=L*, at *t* = 3.15×10^8^ s =10 years. The scenario assumes an infinite half-life of Aβ monomers. The curves marked as “approximate *θ*_1/2,*B*_ →0” in Fig. 5a and S5c are computed utilizing Eqs. (21) and (19), respectively. They are designated as approximate because these equations were obtained using the lumped capacitance approximation. The curve marked as “approximate upper limit” is computed utilizing Eq. (S13). This curve is marked as upper limit because this scenario gives the largest plaque radius (see Fig. 6).

The concentration of free Aβ aggregates, *C*_*B*_, increases from ~0 to ~69 μM (a value predicted by Eq. (S14)) in the range of *θ*_1/2,*B*_ between 10^6^ −10^11^ s (Fig. 5b). A decrease in monomer diffusivity to 6.23 μm^2^/s decreases the maximum value of *C*_*B*_from ~69 μM to ~66 μM (Fig. 5b).

The concentration of Aβ aggregates deposited into senile plaques, *C*_*D*_, starts from a value of ~20 μM at small values of *θ*_1/2,*B*_. This value is predicted by the approximate solution given by Eq. (19) in the case of *θ*_1/2,*B*_→0. Then *C*_*D*_exhibits a rapid increase and then remains at a peak value of ~69 μM in the range of *θ*_1/2,*B*_ between 10^5^ −10^7^ s (Fig. 5c). With further increase in *θ*_1/2,*B*_ the value of *C*_*D*_ decreases to 0, which happens because for *θ*_1/2,*B*_→∞ the free Aβ aggregates do not deposit into the senile plaques (Fig. 5c).

For the scenario with a physiologically relevant half-life of Aβ monomers, *T*_1/ 2, *A*_ = 4.61×10^3^ s, there is still a decrease in *C*_*A*_as the value of *θ*_1/2,*B*_is increased, but the magnitude of this decrease is much smaller (Fig. S8a). The increase in *C*_*B*_when the value of *θ*_1/2,*B*_is increased also occurs, but its magnitude is much smaller (Fig. S8b). The plateau corresponding to the high *C*_*D*_values in Fig. 5c collapses to a single peak value in Fig. S8c.

In the case of an infinitely large half-life of Aβ monomers, the radius of an Aβ plaque undergoes changes between three plateaus (Fig. 6). The first plateau occurs at small values of *θ*_1/2,*B*_and corresponds to *r*_*ABP*_ ≈1.6 μm. The second plateau occurs in the range of *θ*_1/2,*B*_ between approximately 10^5^s and 10^7^s.

**Fig. 6.**
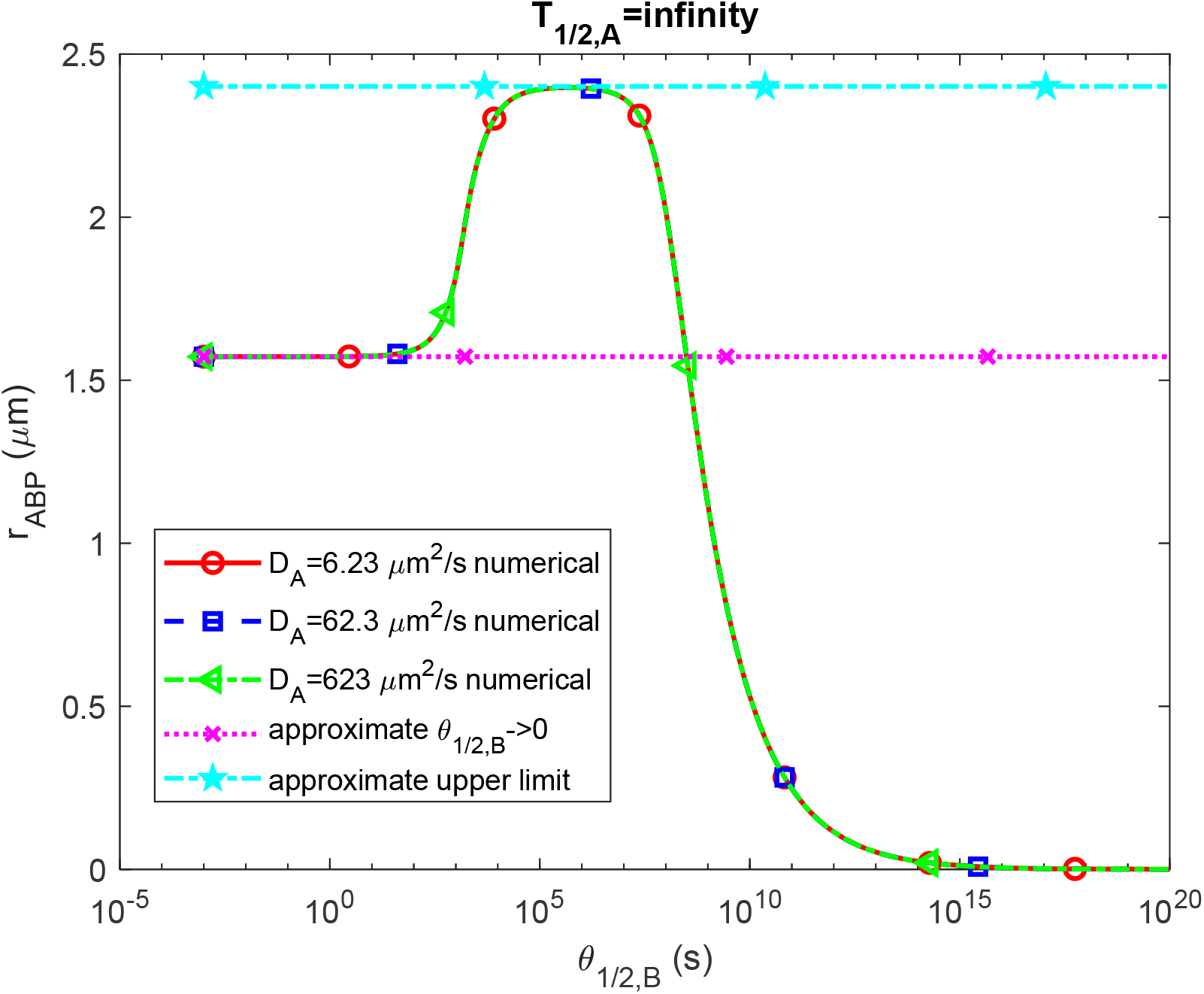
(a) Radius of a growing Aβ plaque, *r*_*ABP*_, vs the half-deposition time of Aβ aggregates into senile plaques, *θ*_1/2,*B*_. The numerical solution is obtained by numerically solving the problem given by Eqs. (3)-(8) and then using Eq. (29) to calculate the radius of the Aβ plaque. The scenario assumes an infinite half-life of Aβ monomers. The curve marked as “approximate *θ*_1/2,*B*_ →0” is computed utilizing Eq. (30). It is designated as approximate because this equation was obtained using the lumped capacitance approximation. The curve marked as “approximate upper limit” is computed utilizing Eq. (S15).

It is the highest plateau, corresponding to *r*_*ABP*_≈ 2.4 μm. The first plateau thus demonstrates the neuroprotective effect of the quick deposition of free Aβ into senile plaques (this corresponds to small values of *θ*_1/2,*B*_). Quick deposition of free aggregates into plaques slows down the rate at which free Aβ aggregates are synthesized and hence leads to a smaller size of Aβ plaques. The ratio of the plaque radius corresponding to the second plateau to that corresponding to the first plateau is 2.4 /1.6 ≈1.5, indicating an approximately 50% reduction in the plaque size. The third plateau occurs when *θ*_1/2,*B*_→0, which corresponds to the situation when free Aβ aggregates do not deposit into plaques. The third plateau corresponds to *r*_*ABP*_= 0 (Fig. 6).

In the case of a physiologically relevant half-life of Aβ monomers, *T*_1/ 2, *A*_ = 4.61×10^3^ s, the second plateau collapses to a peak (Fig. S9). The peak is now much smaller; it corresponds to *r*_*ABP*_≈ 0.08 μm, which is 2.4 / 0.08 ≈ 30 times smaller than in Fig. 6. This means that senile plaques cannot effectively form when the degradation machinery of Aβ monomers functions normally.

In the case of an infinitely large half-life of Aβ monomers, the sensitivity of the radius of the Aβ plaque, *r*_*ABP*_, to the half-deposition time of Aβ aggregates into senile plaques, *θ*_1/2,*B*_, is negative for large values of *θ*_1/2,*B*_ (see, for example, the solid cyan curve corresponding to *θ*_1/2,*B*_ =10^20^ s in Fig. 7a). This implies that an increase in *θ*_1/2,*B*_leads to a decrease in *r*_*ABP*_. This occurs because for large values of *θ*_1/2,*B*_, the deposition of Aβ aggregates into plaques is very slow, resulting in a very small plaque radius (see Fig. 6). A further increase in *θ*_1/2,*B*_reduces the deposition rate even more, resulting in an even smaller plaque radius. The situation is different for small *θ*_1/2,*B*_. In this case, free Aβ aggregates deposit into plaques very quickly. This results in a small concentration of free Aβ aggregates. Since the synthesis of free Aβ aggregates is autocatalytic (see Eq. (2)), an increase in *θ*_1/2,*B*_results in an increase of the rate of synthesis of free Aβ aggregates, and hence in larger *r*. This explains why 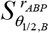 is positive for small values of *θ*_1/2,*B*_.

**Fig. 7.**
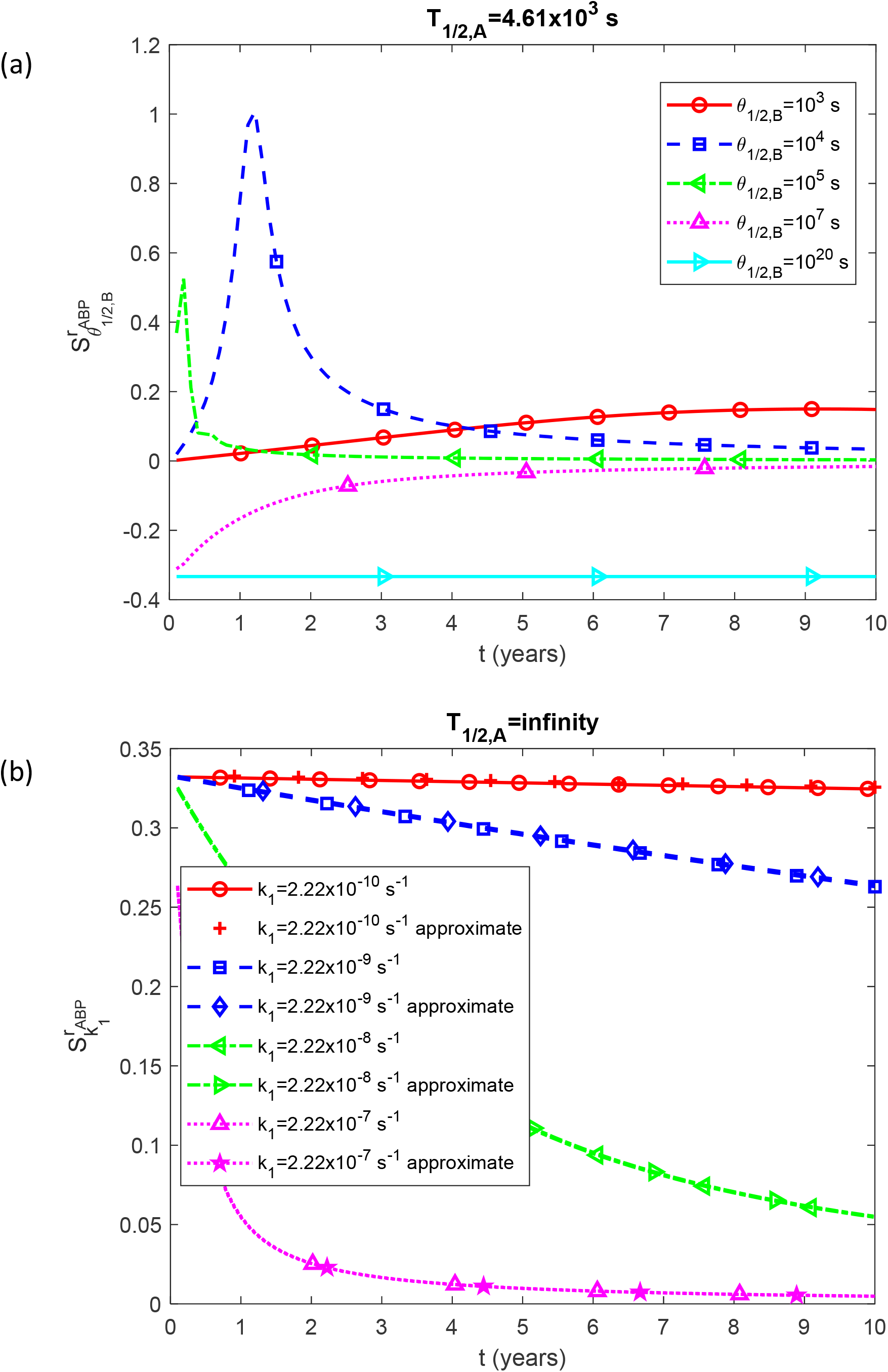
Dimensionless sensitivity of the Aβ plaque’s radius, *r*_*ABP*_, to the following parameters: (a) half-deposition time of Aβ aggregates into senile plaques, *θ*_1/2,*B*_; (b) rate constant that describes nucleation of Aβ aggregates, *k*_1_. The depicted scenario assumes an infinite half-life of Aβ monomers. Fig. 7b is computed for a small half-deposition time of Aβ aggregates into senile plaques, *θ*_1/2,*B*_=10^−3^ s. The curves marked as “approximate” in Fig. 7b are computed utilizing Eq. (36).

The increase in the value of the rate constant describing the nucleation of Aβ aggregates, *k*_1_, leads to an increase in the plaque radius because a larger concentration of free Aβ aggregates results in larger Aβ deposition into plaques (see Eq. (5)). Therefore, 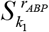 is positive (Fig. 7b). Note the excellent agreement with the approximate solution given by Eq. (36). If *k*_1_ is very small (see the red solid curve in Fig. 7b corresponding to *k*_1_= 2.22 ×10^−10^ s^-1^), 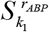 approaches a limiting value of 1/3 given by Eq. (37) (Fig. 7b).

In the case of a physiologically relevant half-life of Aβ monomers, *T*_1/ 2, *A*_ = 4.61×10^3^ s, 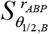 can even change from negative to positive as time increases (see the magenta dotted line in Fig. S10a). This is because initially, the limiting factor is the slow deposition of free Aβ aggregates into plaques. However, for larger times, the limiting factor changes to the small number of Aβ aggregates, resulting in a slow autocatalysis rate. 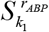 is independent of time and is close to a limiting value of 1/3 (Fig. S10b).

## Discussion, limitations of the model, and future directions

One popular hypothesis associates sporadic forms of AD with impaired clearance of Aβ aggregates (Selkoe, 2001; Tanzi et al., 2004; Saido and Leissring, 2012). The findings from this study suggest that preventing AD does not necessarily require the proteolytic system to degrade Aβ polymers. Instead, the results demonstrate that even if the proteolytic system only targets and degrades Aβ monomers, the aggregation of Aβ cannot progress due to the insufficient supply of reactants (monomers). This result holds significant implications for potential therapeutic interventions in AD.

The effect of the diffusion of Aβ monomers is shown to be small, confirming that the lumped capacitance approximation can be used. The utilization of the lumped capacitance approximation made it possible to obtain approximate solutions that can predict concentrations of Aβ monomers and aggregates and the radius of Aβ plaques for two limiting cases: (i) the case when Aβ aggregates quickly deposit into senile plaques, *θ*_1/2,*B*_→ 0 (section 2.2.2), and (ii) the case of a slow rate of deposition of Aβ aggregates into senile plaques, *θ*_1/2,*B*_→∞ (section S1.2 in Supplementary Materials). The plaque radius for case (ii) is zero (Fig. 6).

Intriguingly, the model predicts that the largest plaque radius occurs not when *θ*_1/2,*B*_→0 or when *θ*_1/2,*B*_→∞, but in the range of *θ* _1/2,*B*_between 10^5^ −10^7^s (Fig. 6). This is explained by the combination of two effects: the rate at which Aβ aggregates deposit into the plaques and the rate of autocatalytic synthesis of free Aβ aggregates. If *θ*_1/2,*B*_→∞, *r*_*ABP*_is equal to zero because free aggregates do not deposit into plaques. However, if *θ*_1/2,*B*_→0, the deposition of free aggregates is also not very fast due to the small concentration of free Aβ aggregates. The small concentration of free aggregates is due to the presence of two mechanisms for the synthesis of free Aβ aggregates: nucleation (see Eq. (1)) and an autocatalytic process (see Eq. (2)). Nucleation is slow, whereas the autocatalytic process is fast (Watzky and Finke, 1997). If free Aβ aggregates are deposited very quickly into the plaques, there are not enough free Aβ aggregates to support the autocatalytic process, resulting in slow synthesis of free Aβ aggregates (primarily by nucleation). This explains the potential neuroprotective effect of senile plaques (Goure et al., 2014). It also elucidates why the largest plaque radius occurs at an intermediate value of *θ*_1/2,*B*_(Fig. 6). It is also interesting that the region where the maximum value of *r*_*ABP*_occurs resembles more a plateau than a sharp peak (Fig. 6).

The current model is limited as the F-W model it relies upon does not simulate the size of Aβ polymers. An enhanced model capable of simulating polymers with varying sizes would significantly enhance the value and accuracy of model predictions. Enhancing the model’s accuracy would be possible with a more precise depiction of the CV’s geometry, particularly focusing on the surface of the lipid membrane where Aβ monomers are generated.

Note that the transformation from the spatially extended model given by Eqs. (3)-(5) to the lumped capacitance model given by Eqs. (9)-(11) can be achieved through volume averaging. If *C*_*A*_, *C*_*B*_, and *C*_*D*_depend solely on *t*, the transformation is straightforward. However, if there is even a slight dependence on *x*, exploring conditions under which non-linear terms can be transformed, such as ⟨*C*_*A*_*C*_*B*_⟩ ≈ ⟨*C*_*A*_⟩⟨*C*_*B*_⟩, is an area of interest for future research.

In future research, it would be important to develop models that simulate the interaction between Aβ aggregates and tau tangles (Hampel et al., 2021). To achieve this, the model would need to account for cross-membrane diffusion processes, as Aβ aggregates are typically formed in the extracellular environment, while tau tangles are intracellular. Additionally, further modeling efforts should consider the activation of microglia in response to Aβ aggregation. It would be crucial to explore the potential combination of microglia-mediated degradation of Aβ aggregates and the neuroinflammatory response triggered by microglia activation (Hansen et al., 2018; Chen et al., 2017; Meyer-Luehmann et al., 2008).

## Abbreviations

Aβ: amyloid beta
AD: Alzheimer’s disease
CV: control volume
F-W: Finke-Watzky

## Data accessibility

This article has no additional data.

## Authors’ contributions

AVK is the sole author of this paper.

## Supplemental Materials

### S1. Dimensionless equations

#### S1.1. Dimensionless form of Eqs. (3)-(8)

The model represented by Eqs. (3)-(8) is reformulated in its dimensionless form as follows:

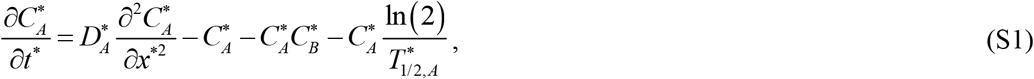

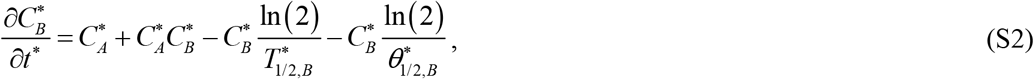

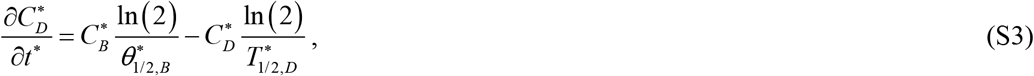

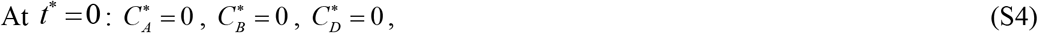

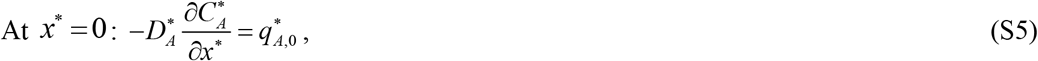

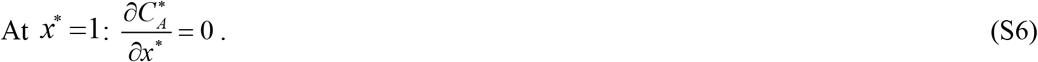

The independent dimensionless variables used in the model are listed in Table S1, and the dependent dimensionless variables are summarized in Table S2. Table S3 gives a summary of the dimensionless parameters utilized in the model.

#### S1.2. Analytical solution of Eqs. (22) and (23) for the scenario where free Aβ aggregates deposit slowly into senile plaques, *θ*_1/2,*B*_→∞

Upon adding Eqs. (22) and (23), the following result is obtained:

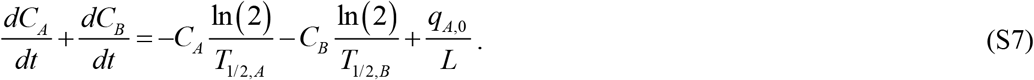

In the case of *T*_1/ 2, *A*_→∞ and *T*_1/ 2, *B*_→∞, integrating Eq. (S7) with respect to time and using the initial condition given by Eq. (6a,b), the following result is obtained:

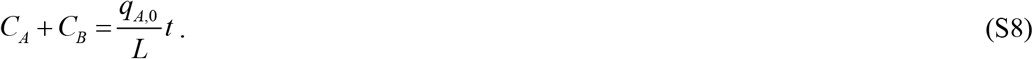

The increase of *C*_*A*_+ *C*_*B*_with time is attributed to the supply of Aβ monomers through the left-hand side boundary of the CV. By eliminating *C*_*A*_from Eq. (23) using Eq. (S8), the following is obtained:

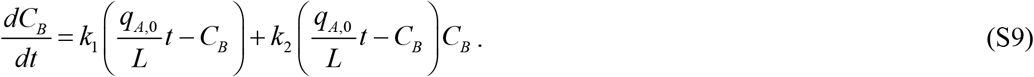

Eq. (S9) is similar to the one studied in Kuznetsov and Kuznetsov (2022). To find the exact solution of Eq. (S9) with the initial condition given by Eq. (6b), the DSolve function followed by the FullSimplify function in Mathematica 13.3 (Wolfram Research, Champaign, IL) was employed. The resulting solution is as follows:

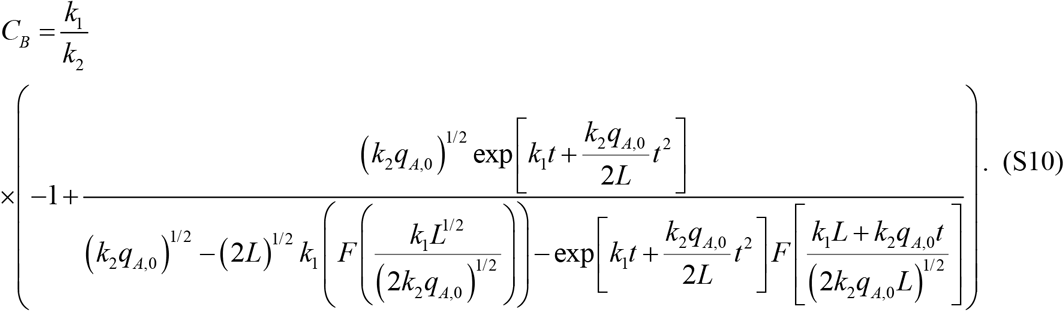

In this equation, *F* (*x*) is Dawson’s integral:

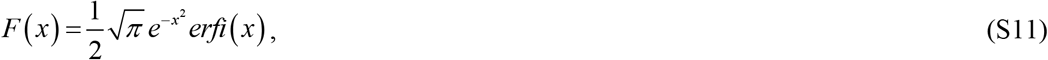

where *erfi* (*x*) is the imaginary error function.

If *t* → 0, Eq. (S10) implies that 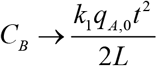, indicating that at small *t*, the concentration of Aβ aggregates is directly proportional to the kinetic constant describing the nucleation of Aβ aggregates. *C*_*A*_can then be found using Eq. (S8) as:

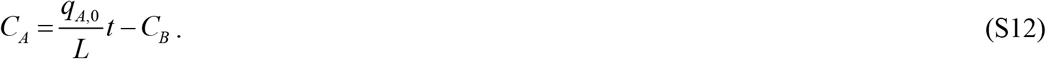

The exact solution provided by Eqs. (S10) and (S12) is quite cumbersome. A more elegant approximate solution, also valid in the case of *T*_1/ 2, *A*_→∞ and *T*_1/ 2, *B*_→∞, is

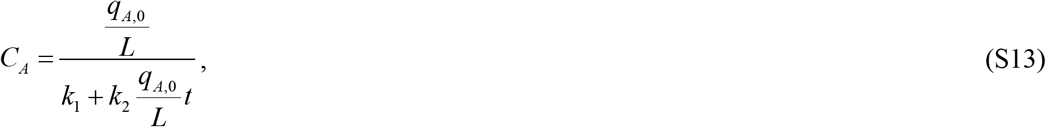

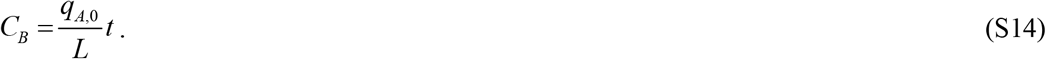

Note that as *t* →∞, Eq. (S13) predicts that *C*_*A*_→ 0.

The upper limit for the plaque radius can be obtained by assuming that all aggregates deposit into plaques. Note that this contradicts the assumption that *θ*_1/2,*B*_→∞, and thus the upper limit can be reached for some finite value of *θ*_1/ 2, *B*_(but not zero). Renaming *C*_*B*_as given in Eq. (S14) to *C*_*D*_and substituting it into Eq. (29), the following is obtained:

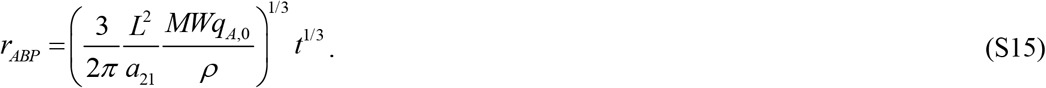

Eq. (S15) represents the cube root of time hypothesis formulated by Kuznetsov (2024a).

#### S1.3. Numerical solution

Eqs. (3)-(5) form a system of two partial differential equations (PDEs). They were solved subject to initial conditions given by Eq. (6) and boundary conditions given by Eqs. (7) and (8) using a well-validated MATLAB PDEPE solver (MATLAB R2020b, MathWorks, Natick, MA, USA). Eqs. (3)-(5) are rewritten in the form required by the PDEPE solver as follows:

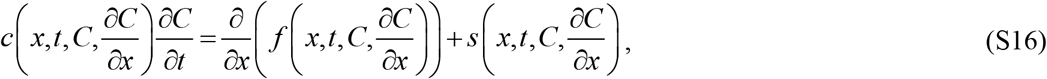

where

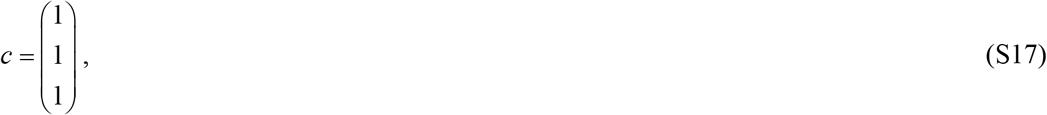

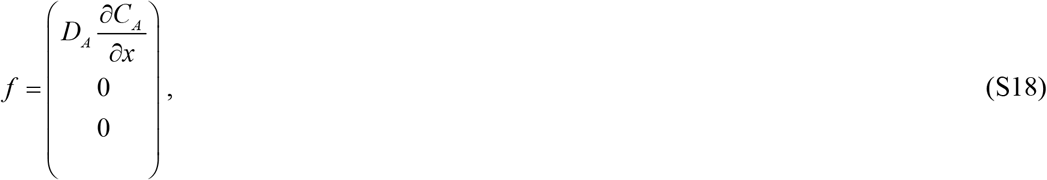

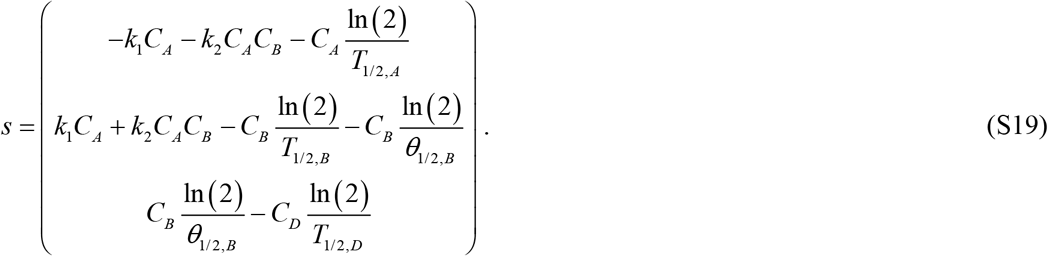

The boundary conditions (7) and (8) are rewritten in the form required by the PDEPE solver as:

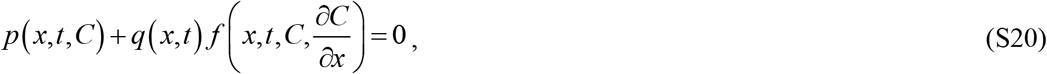

where

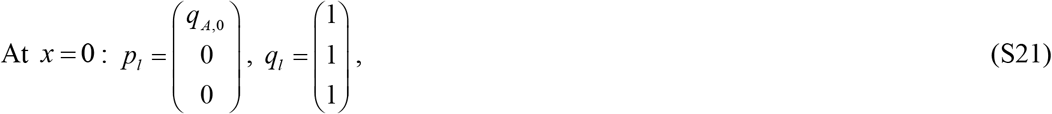

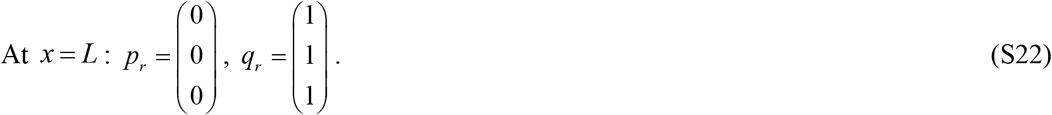

In Eqs. (S21) and (S22), the subscript *l* stands for left, and the subscript *r* stands for right.

Since the second and third elements of vector *f* are equal to zero for any *x* and *t* (Eq. (S18)), setting the second and third components of *f* equal to zero at the boundaries by Eqs. (S21) and (S22) poses no issue.

### S2. Tables

**Table S1.** Dimensionless independent variables used in the model.

| Symbol | Definition |
| --- | --- |
| $t^*$ | $t k_1$ |
| $x^*$ | $x / L$ |

**Table S2.** Dimensionless dependent variables used in the model.

| Symbol | Definition |
| --- | --- |
| $C_A^*$ | $C_A k_2 / k_1$ |
| $C_B^*$ | $C_B k_2 / k_1$ |
| $C_D^*$ | $C_D k_2 / k_1$ |

**Table S3.** Dimensionless parameters used in the model.

| Symbol | Definition | Value(s) obtained using the dimensional parameter values provided in the last column of Table 3 |
| --- | --- | --- |
| $D_A^*$ | $D_A / (L^2 k_1)$ | $1.12 \times 10^7$ |
| $q_{A,0}^*$ | $q_{A,0} k_2 / (k_1^2 L)$ | $4.21 \times 10^5$ |
| $t_f^*$ | $t_f k_1$ | 0.700 |
| $T_{1/2,A}^*$ | $T_{1/2,A} k_1$ | $1.02 \times 10^{-5}$, $2.22 \times 10^{11}$ (to represent $T_{1/2,A}^* \rightarrow \infty$) |
| $T_{1/2,B}^*$ | $T_{1/2,B}k_1$ | $2.22 \times 10^{11}$ (to represent $T_{1/2,B}^* \rightarrow \infty$) |
| $T_{1/2,D}^*$ | $T_{1/2,D}k_1$ | $2.22 \times 10^{11}$ (to represent $T_{1/2,D}^* \rightarrow \infty$) |
| $\theta_{1/2,B}^*$ | $\theta_{1/2,B}k_1$ | $2.22 \times 10^{-12}$, $2.22 \times 10^{-5}$,<br>$2.22 \times 10^{-4}$, $2.22 \times 10^{-2}$,<br>$2.22 \times 10^{11}$ (to represent $\theta_{1/2,B}^* \rightarrow \infty$) |
| $\rho^*$ | $\frac{\rho}{MW} a_{21} \frac{k_2}{k_1}$ | $1.27 \times 10^9$ |

### S3. Supplementary figures

**Fig. S1.**
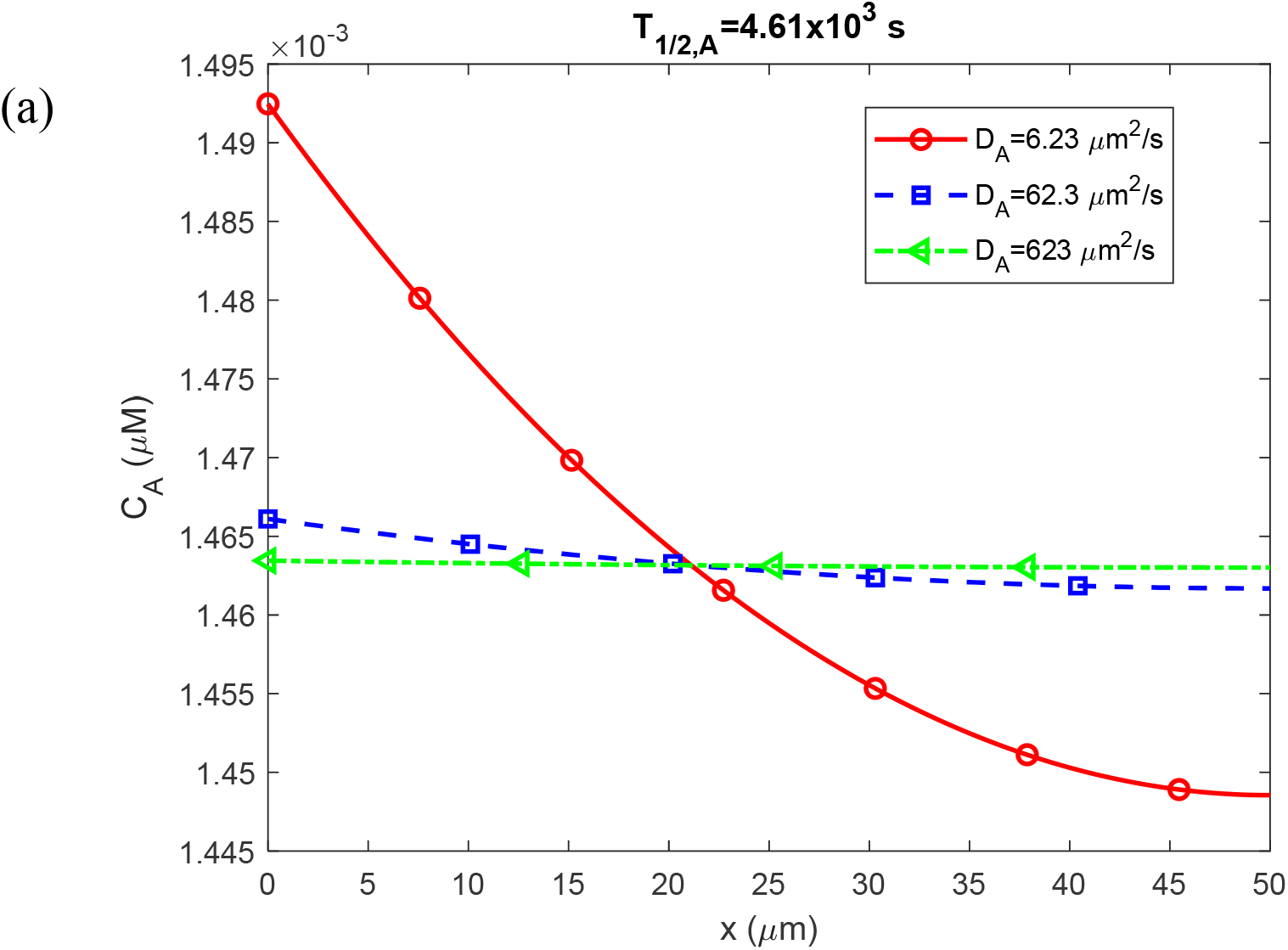
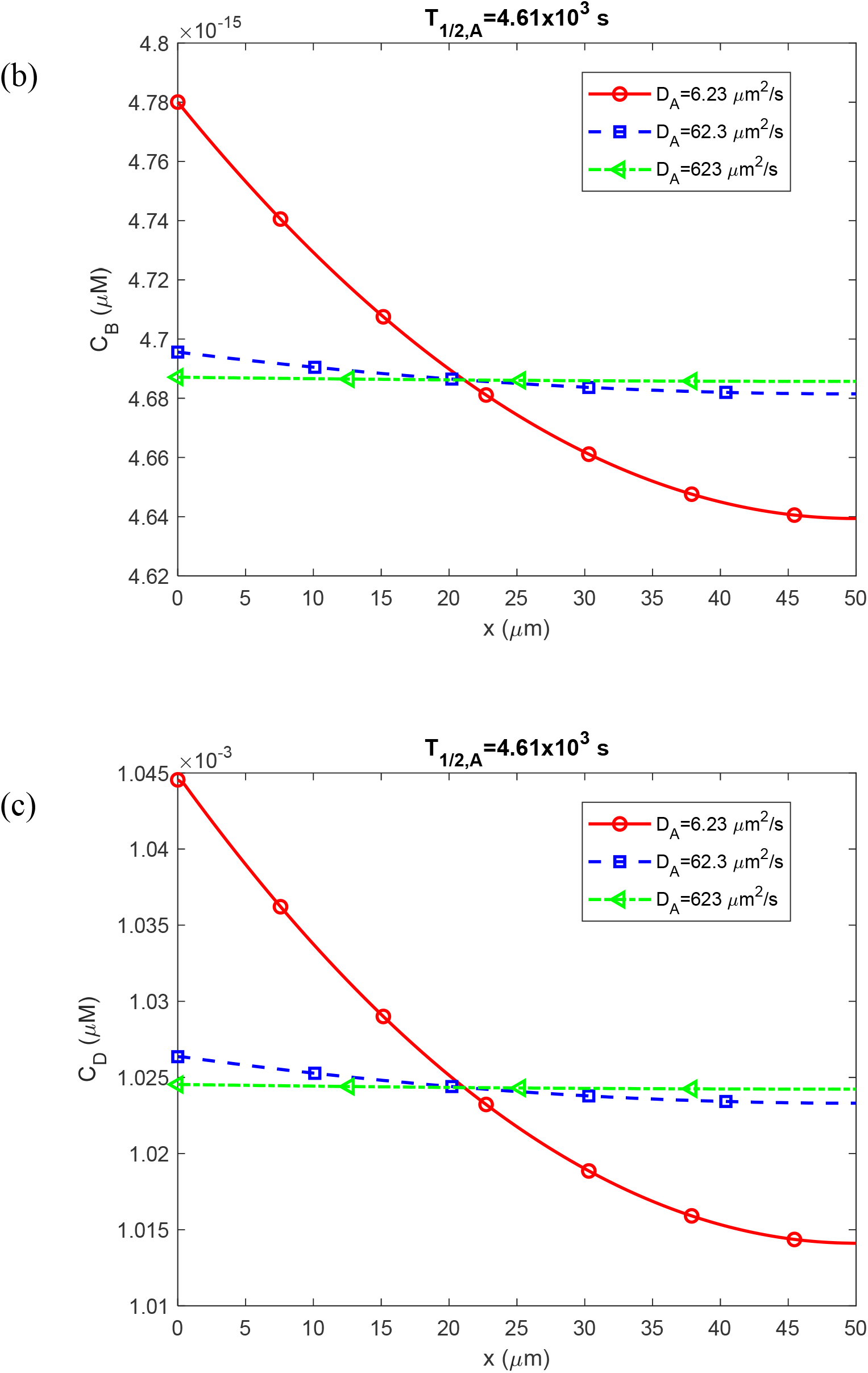
Molar concentration of Aβ monomers, *C*_*A*_(a); free Aβ aggregates, *C*_*B*_(b); and Aβ aggregates deposited into Aβ plaques, *C*_*D*_(c); as a function of the distance from the surface that releases Aβ monomers (e.g., cell membrane). These computational results are presented at *t* = 3.15×10^8^ s =10 years and correspond to the scenario with a finite half-life of Aβ monomers, *T*_1/ 2, *A*_ = 4.61×10^3^ s, and a small half-deposition time of Aβ aggregates into senile plaques, *θ*_1/2,*B*_=10^−3^ s.

**Fig. S2.**
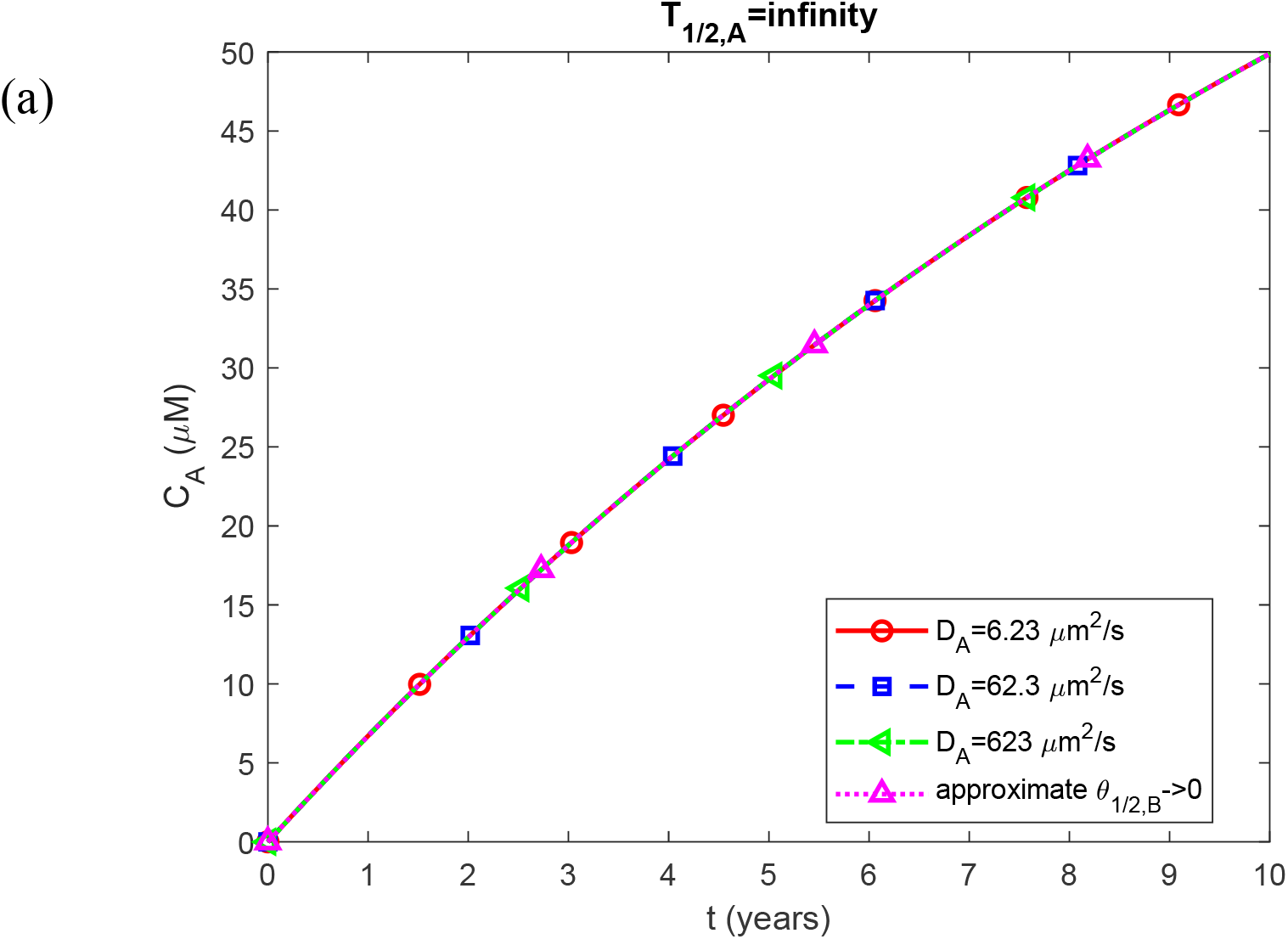
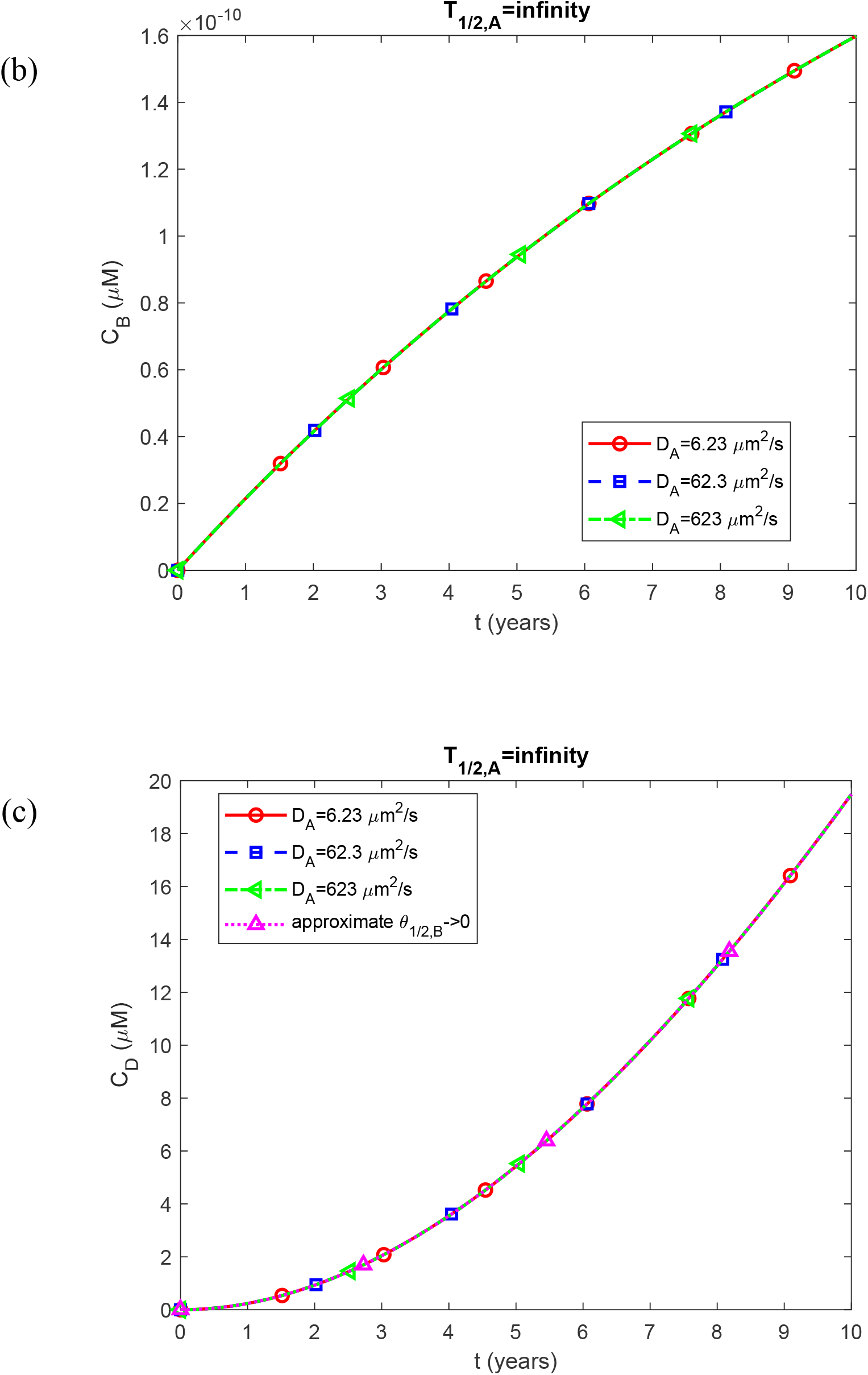
Molar concentration of Aβ monomers, *C*_*A*_(a); free Aβ aggregates, *C*_*B*_(b); and Aβ aggregates deposited into Aβ plaques, *C*_*D*_(c); as a function of time (in years). The computational results are presented at the right-hand side boundary of the CV, *x* = *L*. The scenario displayed in this figure assumes an infinite half-life of Aβ monomers and a small half-deposition time of Aβ aggregates into senile plaques, *θ*_1/2,*B*_ =10^−3^ s. The curves marked as “approximate *θ*_1/2,*B*_ →0” in Fig. S2a and S2c are computed utilizing Eqs. (21) and (19), respectively. They are designated as approximate because these equations were obtained using the lumped capacitance approximation.

**Fig. S3.**
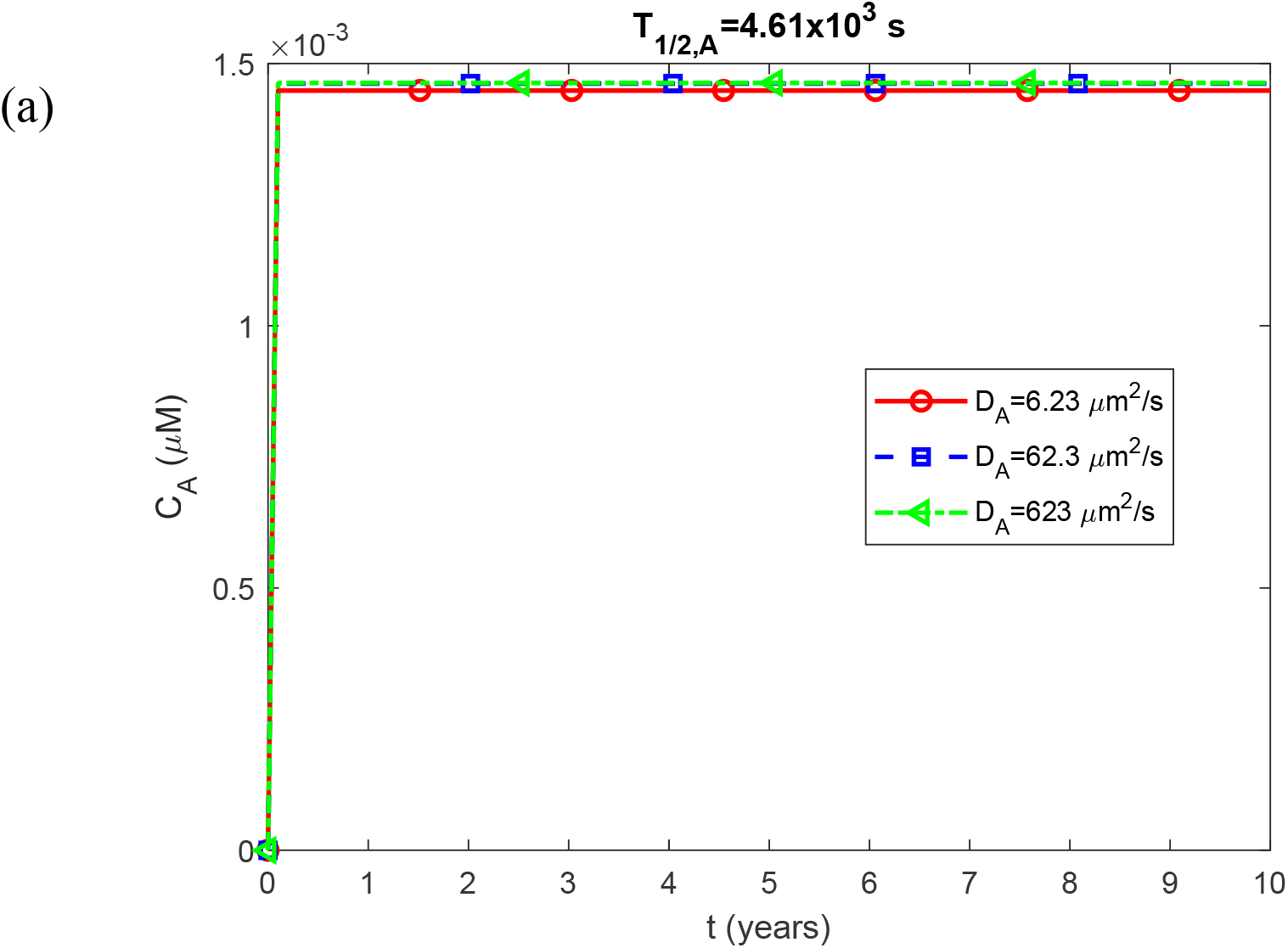
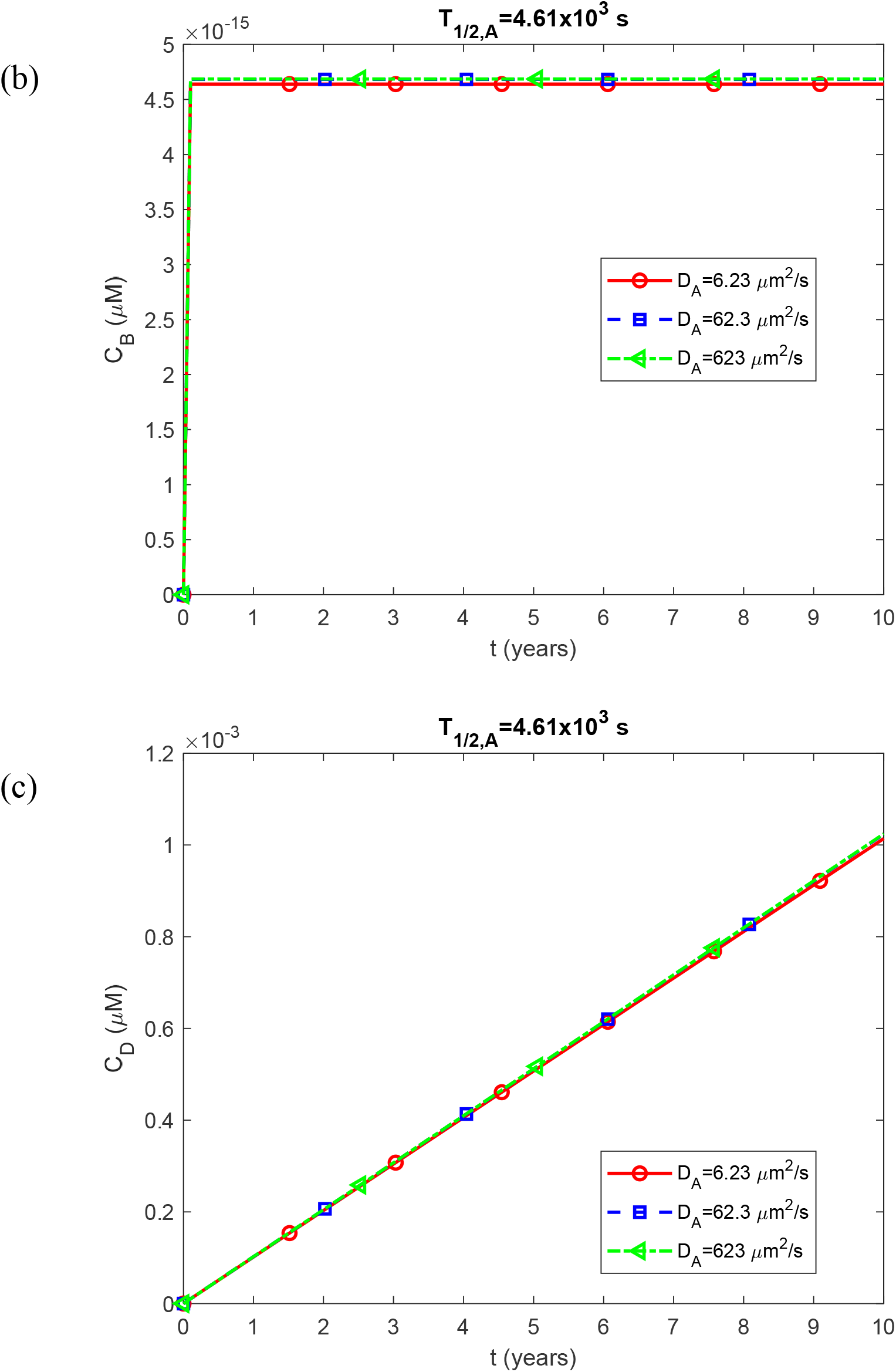
Molar concentration of Aβ monomers, *C*_*A*_(a); free Aβ aggregates, *C*_*B*_(b); and Aβ aggregates deposited into Aβ plaques, *C*_*D*_(c); as a function of time (in years). The computational results are presented at the right-hand side boundary of the CV, *x* = *L*. The scenario displayed in this figure assumes a finite half-life of Aβ monomers, *T*_1/ 2, *A*_ = 4.61×10^3^ s, and a small half-deposition time of Aβ aggregates into senile plaques, *θ*_1/2,*B*_=10^−3^ s.

**Fig. S4.**
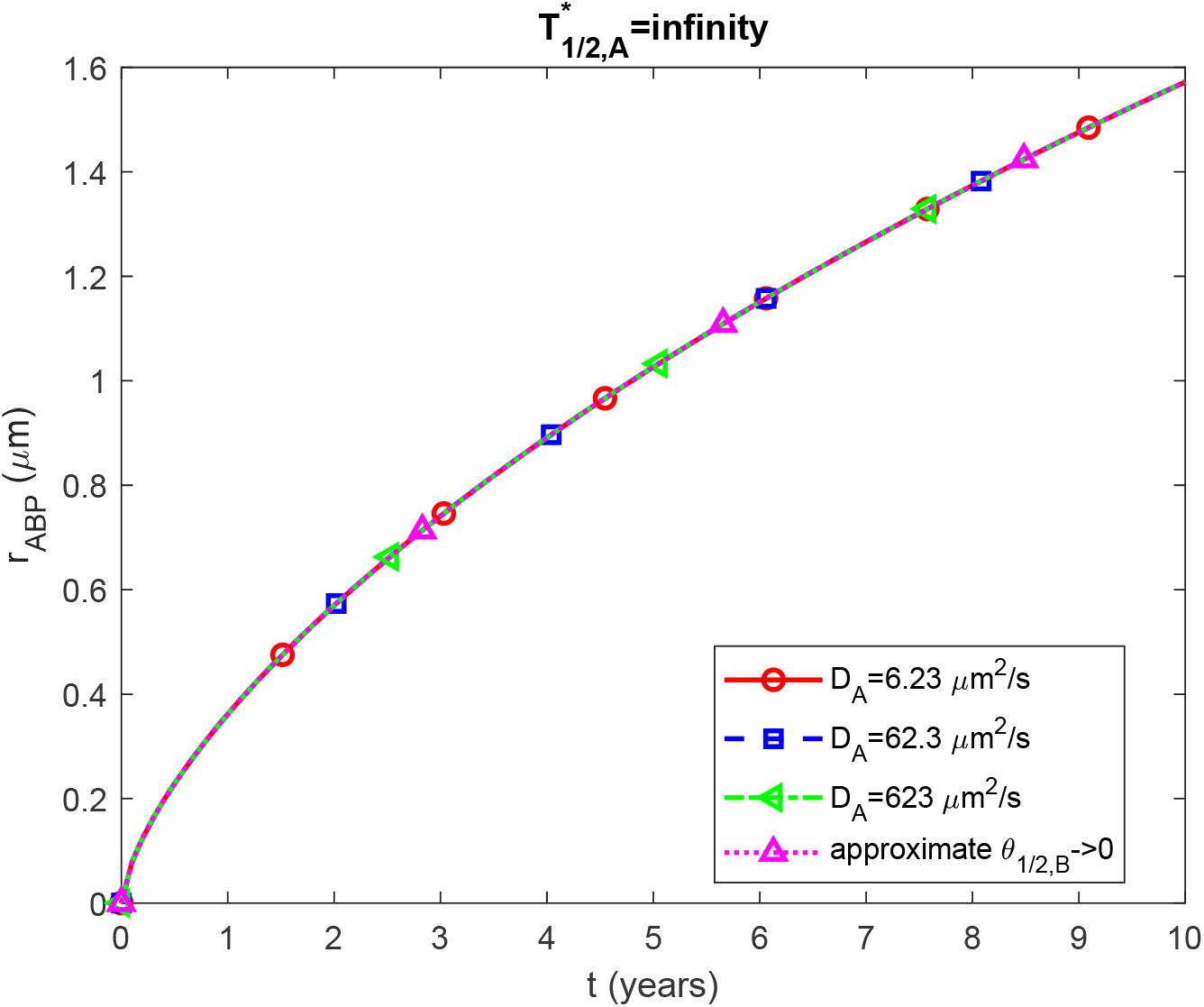
(a) Radius of a growing Aβ plaque, *r*_*ABP*_, vs time (in years). The numerical solution is obtained by numerically solving the problem given by Eqs. (3)-(8) and then using Eq. (29) to calculate the radius of the Aβ plaque. The scenario displayed in this figure assumes an infinite half-life of Aβ monomers and a small half-deposition time of Aβ aggregates into senile plaques, *θ*_1/2,*B*_=10^−3^ s. The curve marked as “approximate *θ*_1/2,*B*_ →0” is computed utilizing Eq. (30). It is designated as approximate because this equation was obtained using the lumped capacitance approximation.

**Fig. S5.**
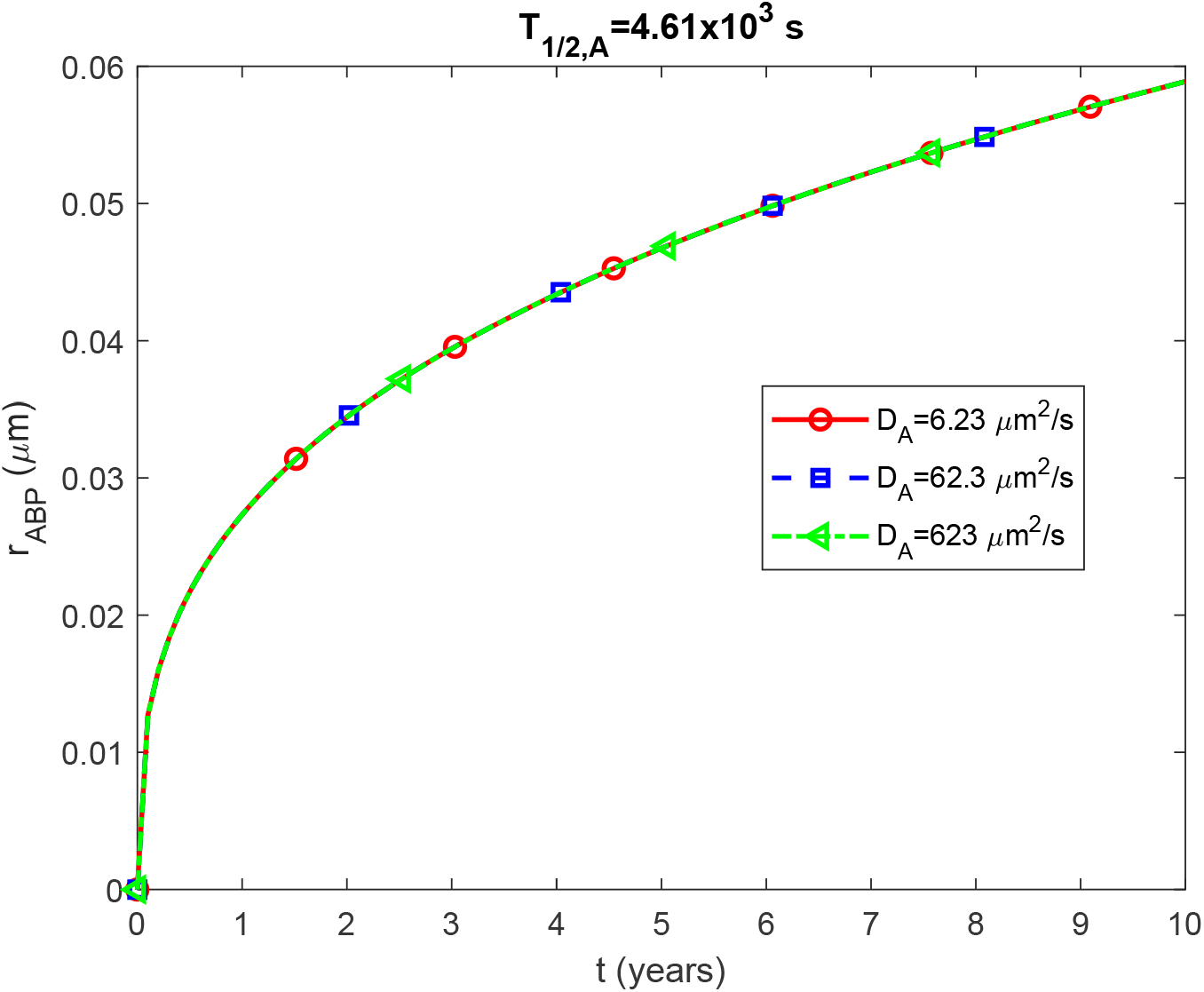
(a) Radius of a growing Aβ plaque, *r*_*ABP*_, vs time (in years). The numerical solution is obtained by numerically solving the problem given by Eqs. (3)-(8) and then using Eq. (29) to calculate the radius of the Aβ plaque. The scenario displayed in this figure assumes a finite half-life of Aβ monomers, *T*_1/ 2, *A*_ = 4.61×10^3^ s, and a small half-deposition time of Aβ aggregates into senile plaques, *θ*_1/2,*B*_ =10^−3^ s.

**Fig. S6.**
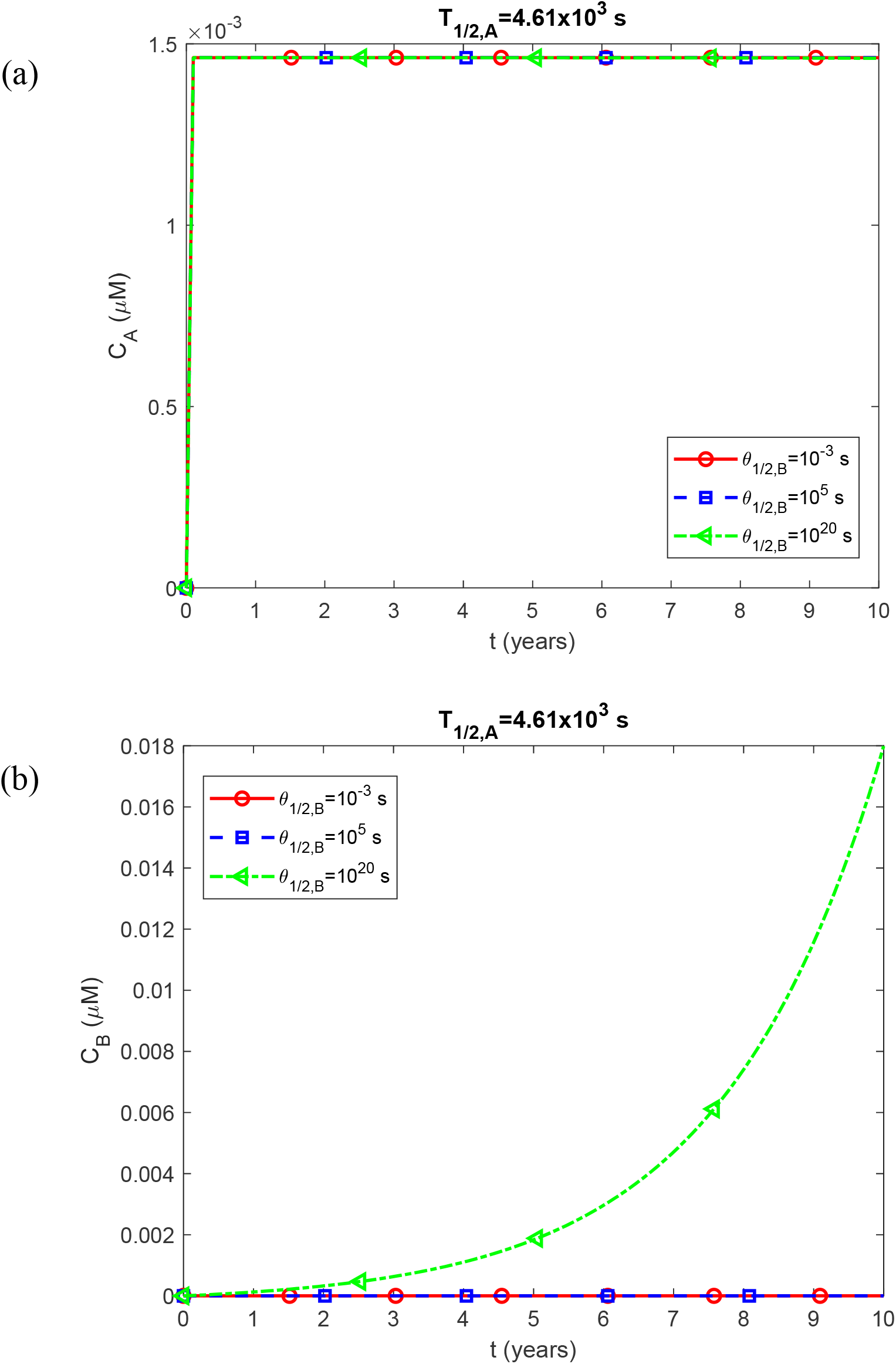
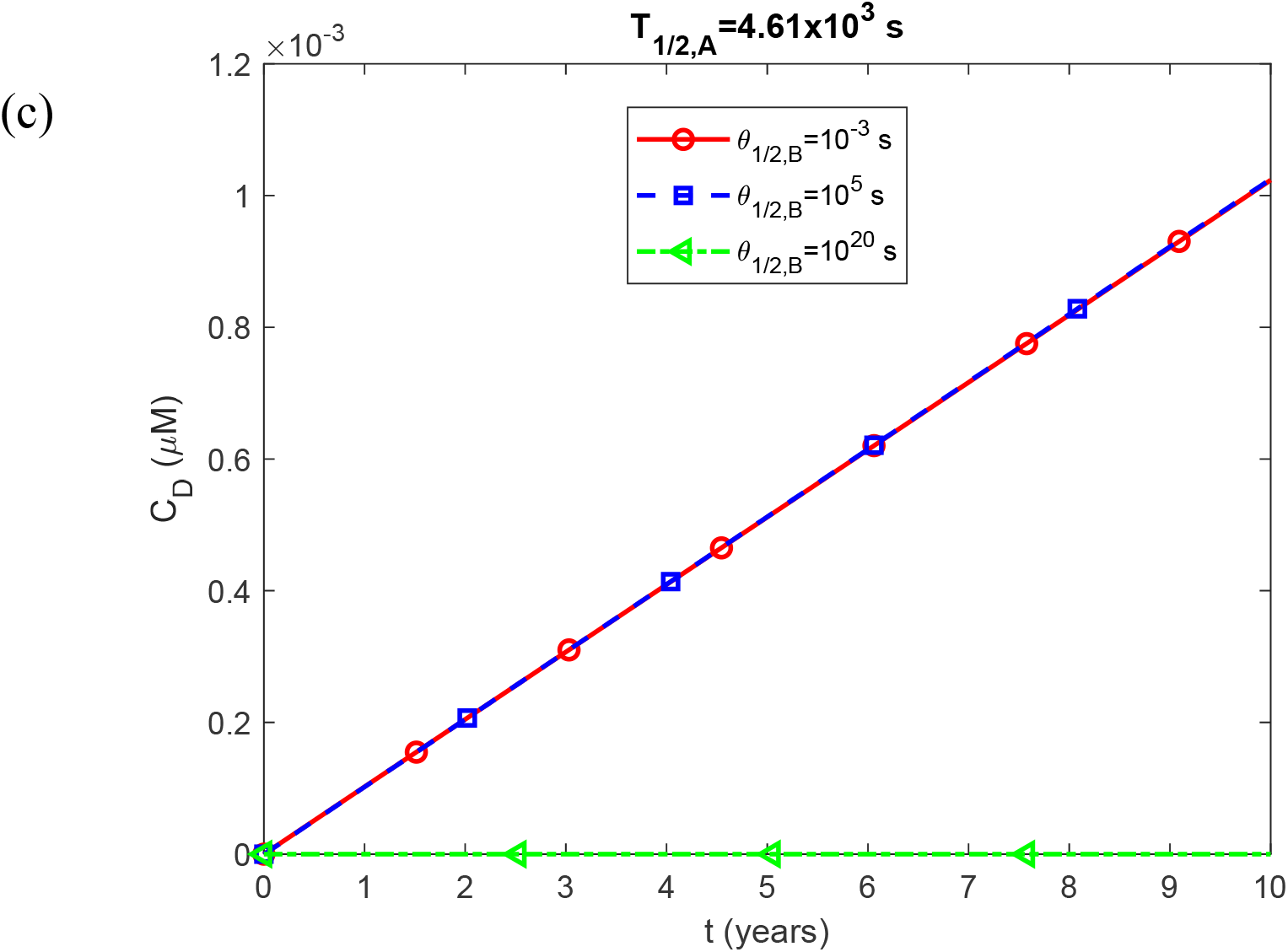
(a) Molar concentration of Aβ monomers, *C*_*A*_(a); free Aβ aggregates, *C*_*B*_(b); and Aβ aggregates deposited into Aβ plaques, *C*_*D*_(c); as a function of the half-deposition time of Aβ aggregates into senile plaques, *θ*_1/2,*B*_. The computational results are presented at the right-hand side boundary of the CV, *x=L*, at *t* = 3.15×10^8^ s =10 years. The scenario assumes a finite half-life of Aβ monomers, *T*_1/ 2, *A*_ = 4.61×10^3^ s.

**Fig. S7.**
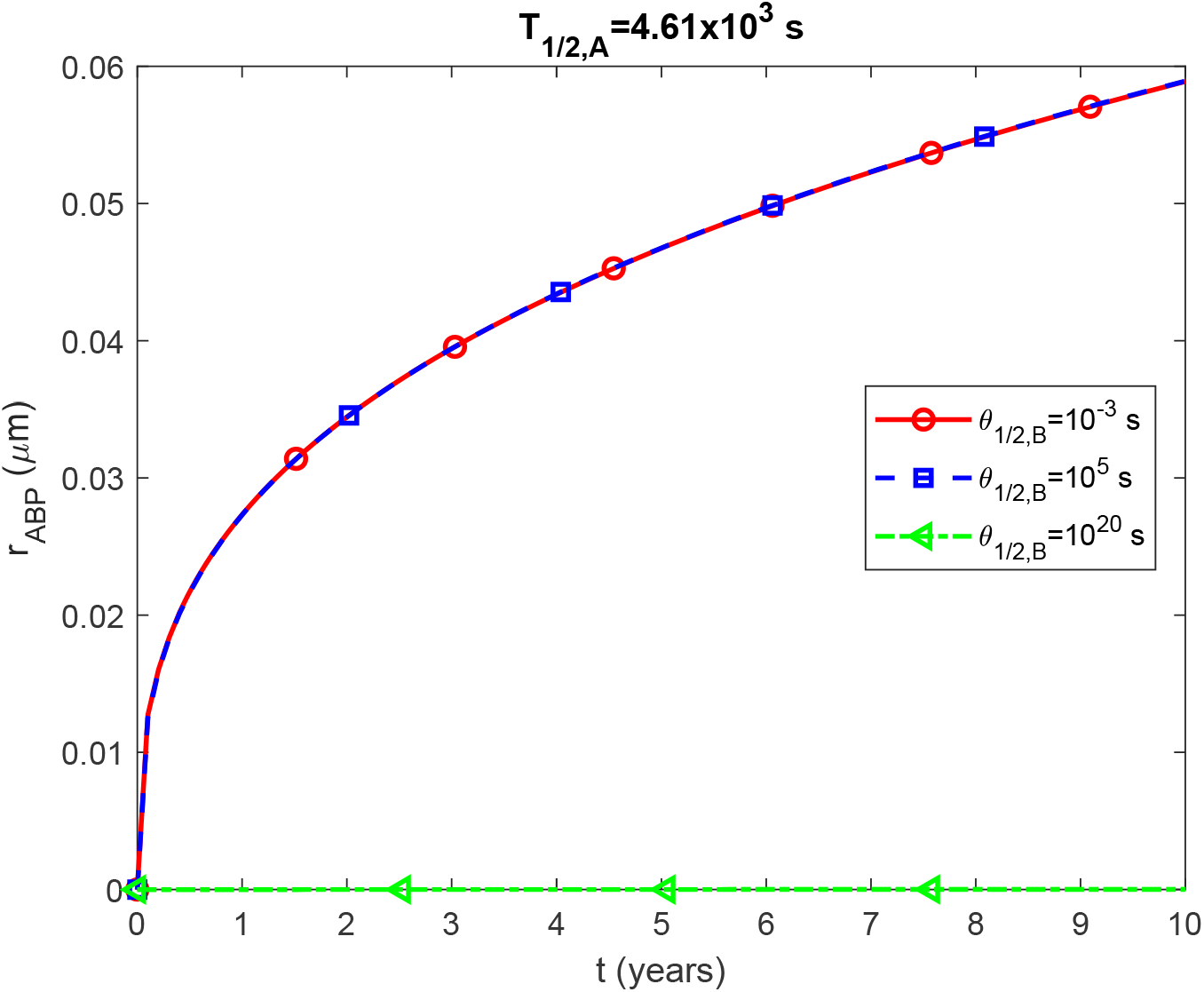
(a) Radius of a growing Aβ plaque, *r*_*ABP*_, vs time (in years). The numerical solution is obtained by numerically solving the problem given by Eqs. (3)-(8) and then using Eq. (29) to calculate the radius of the Aβ plaque. The scenario assumes a finite half-life of Aβ monomers, *T*_1/ 2, *A*_ = 4.61×10^3^ s.

**Fig. S8.**
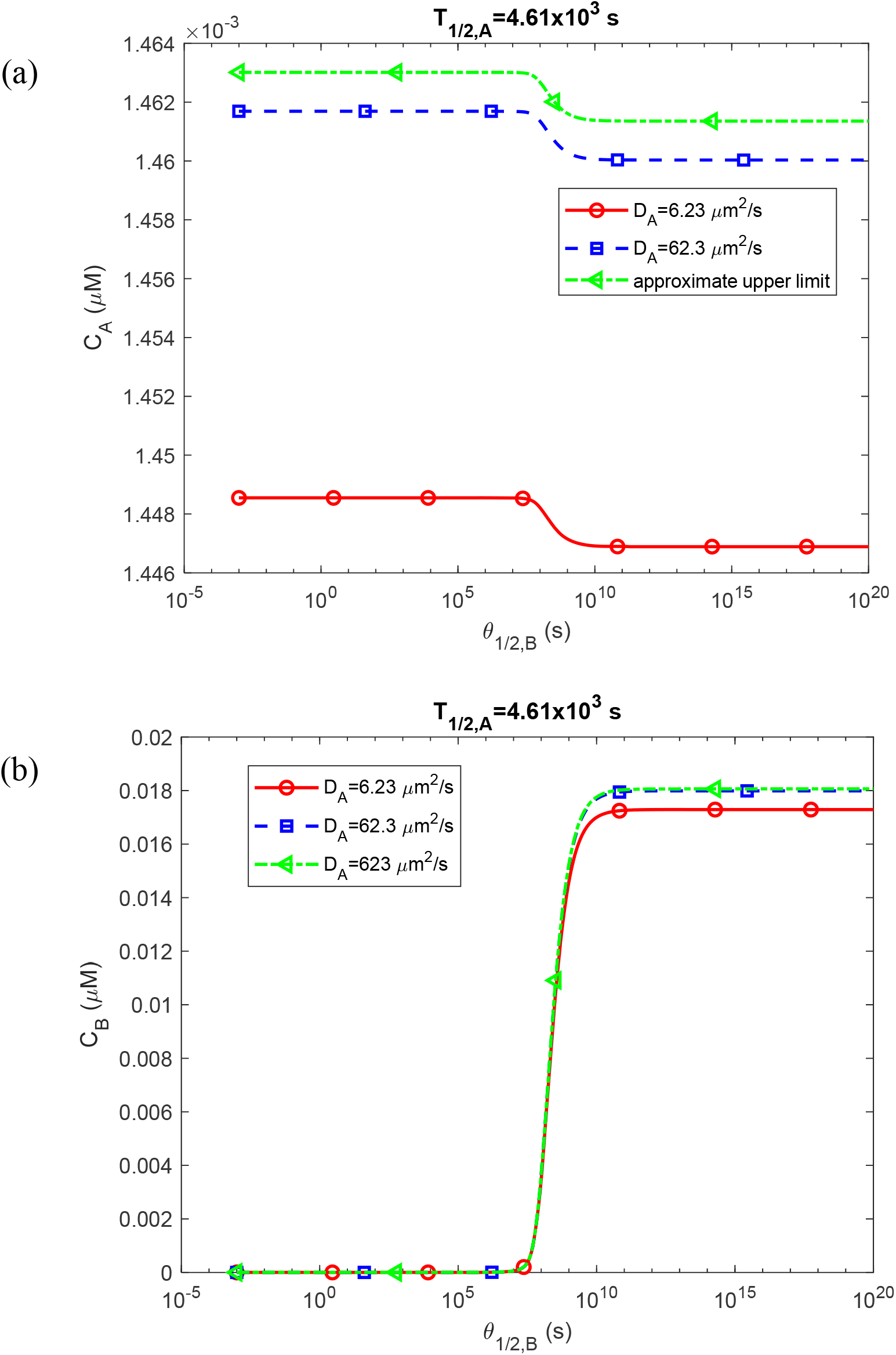
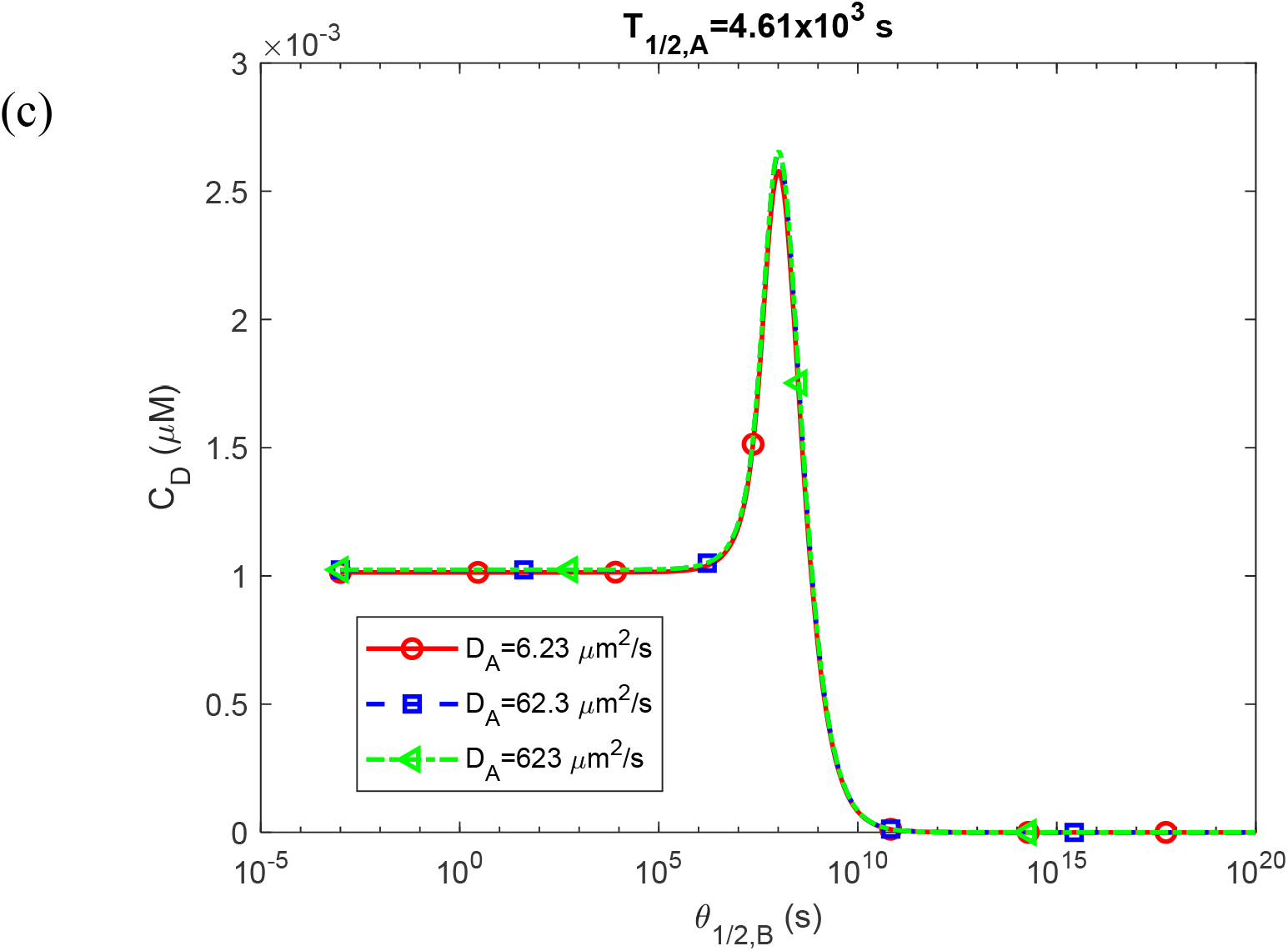
Molar concentration of Aβ monomers, *C*_*A*_(a); free Aβ aggregates, *C*_*B*_(b); and Aβ aggregates deposited into Aβ plaques, *C*_*D*_(c); as a function of the half-deposition time of Aβ aggregates into senile plaques, *θ*_1/2,*B*_. The computational results are presented at the right-hand side boundary of the CV, *x=L*, at *t* = 3.15×10^8^ s =10 years. The scenario assumes a finite half-life of Aβ monomers, *T*_1/ 2, *A*_ = 4.61×10^3^ s.

**Fig. S9.**
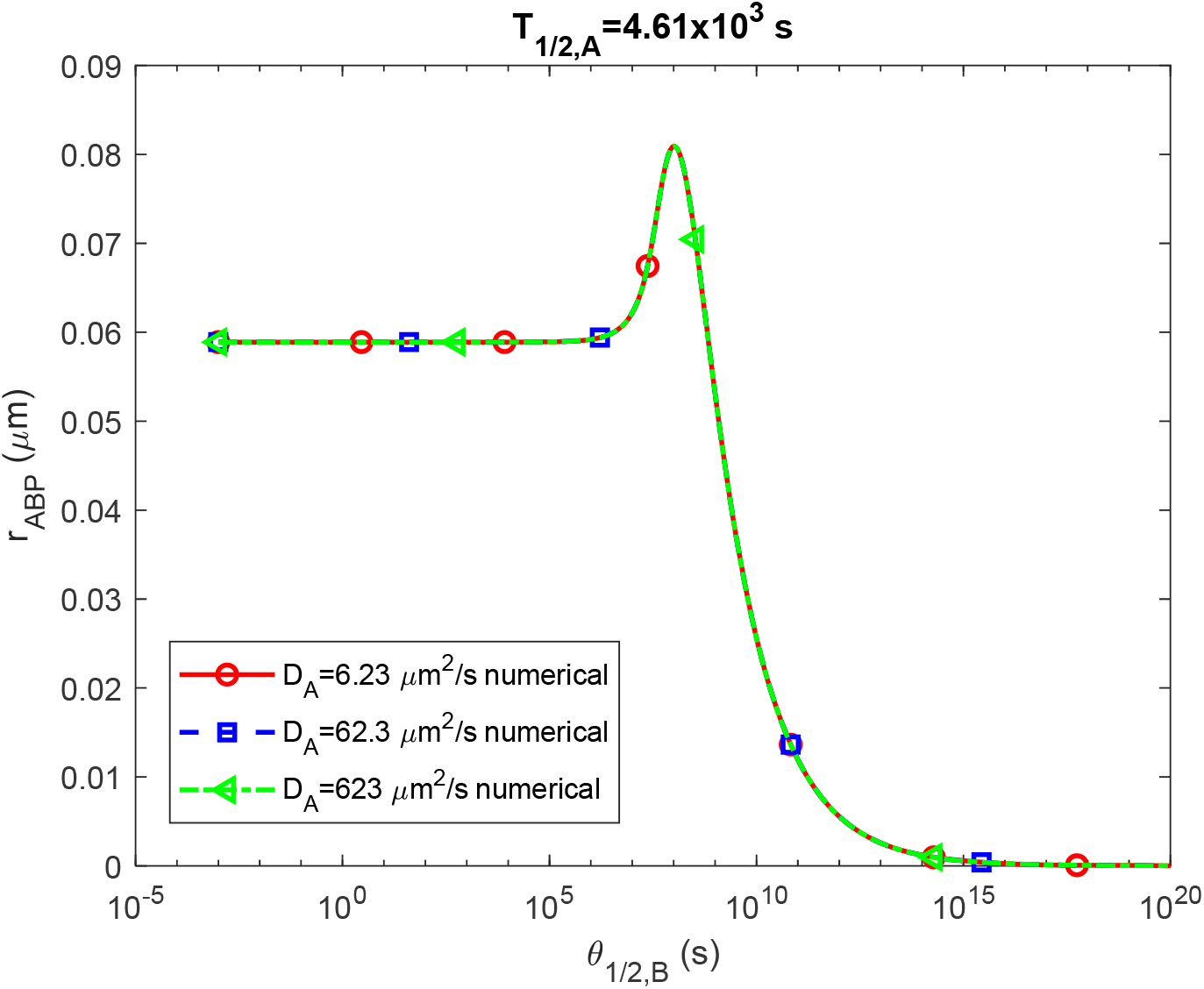
(a) Radius of a growing Aβ plaque, *r*_*ABP*_, vs the half-deposition time of Aβ aggregates into senile plaques, *θ*_1/2,*B*_. The numerical solution is obtained by numerically solving the problem given by Eqs. (3)-(8) and then using Eq. (29) to calculate the radius of the Aβ plaque. The scenario assumes a finite half-life of Aβ monomers, *T*_1/ 2, *A*_ = 4.61×10^3^ s.

**Fig. S10.**
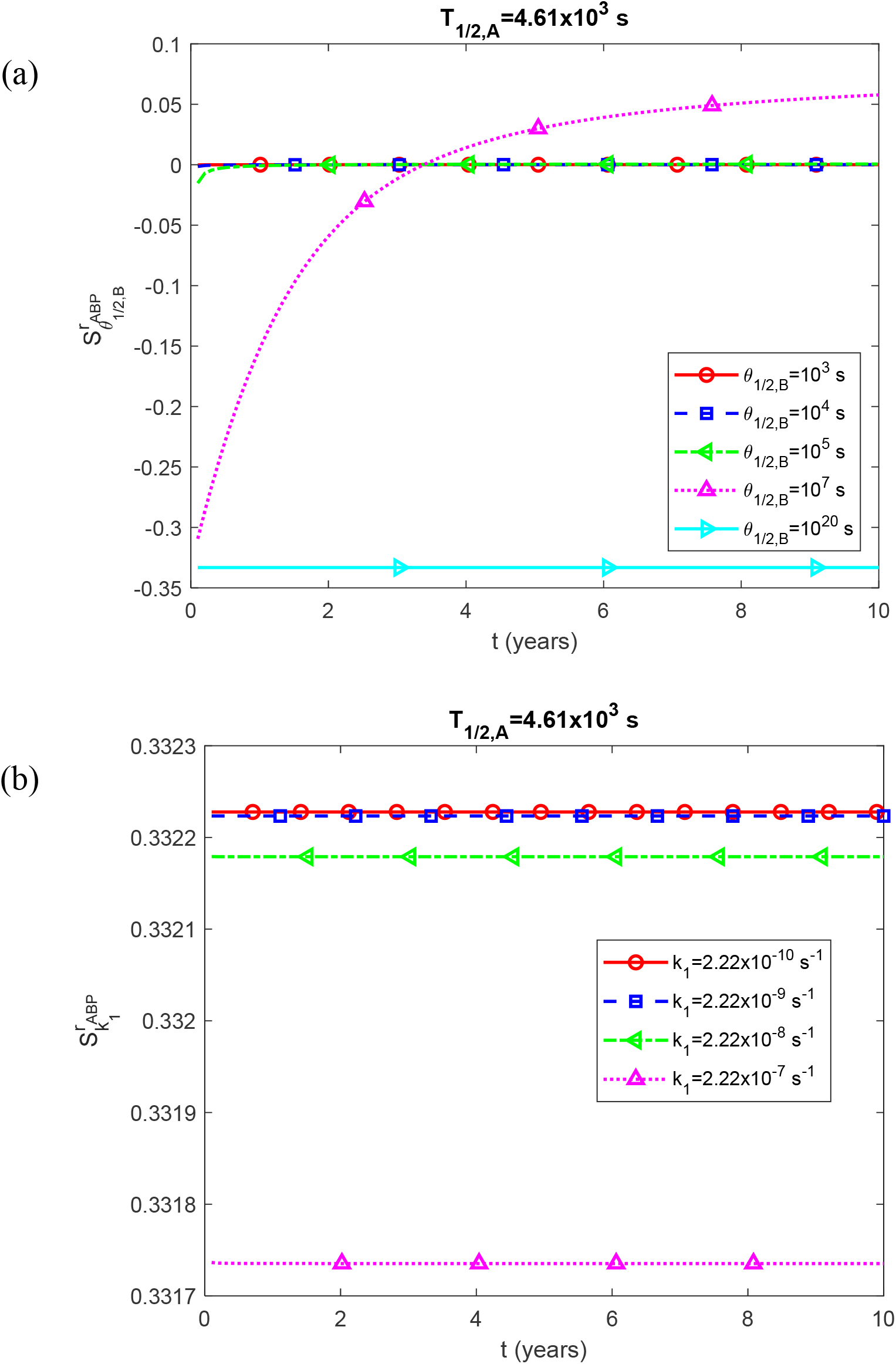
Dimensionless sensitivity of the Aβ plaque’s radius, *r*_*ABP*_, to the following parameters: (a) half-deposition time of Aβ aggregates into senile plaques, *θ*_1/2,*B*_; (b) rate constant that describes nucleation of Aβ aggregates, *k*_1_. The scenario assumes a finite half-life of Aβ monomers, *T*_1/ 2, *A*_ = 4.61×10^3^ s. Fig. S10b is computed for a small half-deposition time of Aβ aggregates into senile plaques, *θ*_1/2,*B*_=10^−3^ s.

